# Saving the Dinaric lynx: multidisciplinary monitoring and stakeholder engagement support large carnivore restoration in human-dominated landscape

**DOI:** 10.1101/2024.10.15.617164

**Authors:** Miha Krofel, Urša Fležar, Rok Černe, Lan Hočevar, Marjeta Konec, Aleksandra Majić Skrbinšek, Tomaž Skrbinšek, Seth Wilson, Bernarda Bele, Jaka Črtalič, Tomislav Gomerčić, Tilen Hvala, Jakub Kubala, Pavel Kvapil, Meta Mavec, Anja Molinari-Jobin, Paolo Molinari, Elena Pazhenkova, Hubert Potočnik, Teodora Sin, Magda Sindičić, Ira Topličanec, Teresa Oliveira

## Abstract

Translocations are central to large carnivore restoration efforts, but inadequate monitoring often inhibits effective conservation decision-making. Extinctions, reintroductions, poaching and high inbreeding levels of the Central European populations of Eurasian lynx (*Lynx lynx*) typify the carnivore conservation challenges in the Anthropocene. Recently, several conservation efforts were initiated to improve the genetic and demographic status, but were met with variable success. Here, we report on a successful, stakeholder-engaged translocation effort to reinforce the highly-inbred Dinaric lynx population and create a new stepping-stone subpopulation in the Southeastern Alps. We used multidisciplinary and internationally-coordinated monitoring using systematic camera-trapping, non-invasive genetic sampling, GPS-tracking of translocated and remnant individuals, recording of reproductive events and interspecific interactions, as well as the simultaneous tracking of the public and stakeholders’ support of carnivore conservation before, during and after the translocation process across the three countries. Among the 22 translocated wild-caught Carpathian lynx, 68% successfully integrated into the population and local ecosystems and at least 59% reproduced. Probability of dispersing from the release areas was 3-times lower when soft-release rather than hard-release method was used. Translocated individuals had lower natural mortality, higher reproductive success and similar ungulate kill rates compared to the remnant lynx. Cooperation with local hunters and protected area managers enabled us to conduct multi-year camera-trapping and non-invasive genetic monitoring across a 12,000-km^2^ transboundary area. Results indicate a reversal in population decline, as the lynx abundance increased for >40% during the 4-year translocation period. Effective inbreeding decreased from 0.32 to 0.08-0.19, suggesting a 2- to 4-fold increase in fitness. Furthermore, successful establishment of a new stepping-stone subpopulation represents an important step towards restoring the Central European lynx metapopulation. Robust partnerships with local communities and hunters coupled with transparent communication helped maintain high public and stakeholder support for lynx conservation throughout the translocation process. Lessons learned about the importance of stakeholder involvement and multidisciplinary monitoring conducted across several countries provide a successful example for further efforts to restore large carnivores in human-dominated ecosystems.

## 1. Introduction

Large carnivores can play a key role in the structure and function of ecosystems (Ripple et al. 2014). However, they are disproportionately affected by humans due to their large habitat needs, apex trophic position, frequent conflicts with people and high levels of human-caused mortality (Darimont et al. 2015; Krofel et al. 2015; Ripple et al. 2014). To counteract declines of large carnivore populations and restore functional ecosystems, conservationists and managers increasingly use translocations to reintroduce extirpated species to their historic ranges or to reinforce declining populations (Thomas et al. 2023). Although survivorship of translocated large carnivores has improved in the last decades and it is higher compared to other vertebrates, only 66% of individuals survive the first six months and 37% reproduce (Thomas et al. 2023). This necessitates better and more holistic understanding of the translocation processes to enable evidence-based and adaptive decision-making (Taylor et al. 2017).

Successful large carnivore translocation efforts must consider both biological and social factors that are context-specific to the species, ecosystems, cultural context, and landscapes where large carnivore recovery efforts take place. These include choosing optimal individual animals and release sites (Thomas et al. 2023), assessing the likelihood of successful integration into local ecosystems (e.g. predation and other interspecific interactions; Hayward and Somers 2009), as well as understanding public attitudes and support among key stakeholder groups (Wilson 2018). Many past monitoring efforts have been limited to tracking movement of translocated animals, and have paid less attention to the broader population impacts or how the translocation process influenced the public and stakeholder attitudes (Taylor et al. 2017; Wilson 2018). Large carnivores are among the most conflict-prone species, and their translocations often result in controversies and public opposition, making it as much a political as a biological challenge (Treves and Karanth 2003). This makes understanding public attitudes an essential part of any reinforcement or reintroduction process and is especially relevant in human-dominated landscapes, like Central Europe, where the scale of large carnivore populations transcends national borders, cultures, and management jurisdictions (Penteriani et al. 2018).

The Eurasian lynx (*Lynx lynx*; hereafter lynx) is an apex predator specialized in hunting wild ungulates and is considered a species of conservation concern in Europe (von Arx et al. 2021). While large, autochthonous populations persist in Northern and Eastern Europe, lynx were exterminated from Western and Central Europe by the beginning of the 20^th^ century. Since the 1970s, several reintroduction attempts have been made, some of which restored lynx to portions of their historic ranges (Linnell et al. 2009). However, these reintroduced populations remain isolated and are currently all classified as endangered or critically endangered (von Arx et al. 2021). These populations are primarily threatened by inbreeding depression (Mueller et al. 2022) and poaching resulting from low acceptance by certain members of local hunting communities that perceive lynx as competitors for wild ungulates, as well as due to other, deeper-rooted motives (Heurich et al. 2018). Recently, several conservation efforts were initiated to improve the genetic and demographic status of these lynx populations through reinforcement of the existing populations or creating new stepping stones to promote gene flow and resilience (Molinari et al. 2021; Port et al. 2024).

The Dinaric lynx population was considered the most inbred lynx population in Europe (Mueller et al. 2022; Sindičić et al. 2013). This population originated from the 1973 translocation of six individuals (some of which were related to one another) from the Carpathian population to the Dinaric Mountains of Slovenia. Although initially the effort was successful as lynx reproduced and expanded their population to neighboring countries along the Dinaric Mountains (Croatia and Bosnia and Herzegovina) and the Southeastern Alps (Italy and Austria), by the 1990s the population expansion stalled and rapidly declined after the 2000s (Fležar et al. 2021). By 2019, lynx were functionally extinct in the Southeastern Alps and the remnant population in the Dinaric mountains was highly inbred (Fe > 0.3), showed signs of inbreeding depression, and faced immediate extinction risk (Fležar et al. 2021; Molinari et al. 2021; Sindičić et al. 2013).

Following favorable public attitude surveys supportive of lynx conservation (Majić Skrbinšek 2008), several international workshops were convened by leading experts to develop a reintroduction plan in 2010s (Breitenmoser 2011; Krofel et al. 2010). Based on an agreement that wild-caught lynx from the Carpathian population would be the most suitable source of lynx for the reinforcement project, the European Union (EU)-funded LIFE Lynx project started in 2017. During the first two years of the project, LIFE Lynx project team members in Slovenia, Croatia and Italy engaged in planning, outreach, education, and meaningful engagement with the public, local communities, and hunters to build support for the translocations (for details see the Supplementary Methods, Section 1.5.1). This was followed by the translocation of 18 lynx from the Carpathian population (Romania and Slovakia) to the Dinaric Mountains and Southeastern Alps in Slovenia and Croatia in 2019-2023. An additional four lynx were translocated from the Carpathian and Jura populations to the Southeastern Alps in Italy in 2023 in the frame of the Urgent Lynx Conservation Action (ULyCA2) project.

These two projects shared three main goals: 1) use population reinforcement to prevent the extinction of the Dinaric population, 2) reintroduce lynx to the Julian Alps to create a new stepping-stone subpopulation in the Southeastern Alps to facilitate connection of the Dinaric and (Western) Alpine populations, and 3) maintain support for lynx conservation among the public and key stakeholders (hunters) through strategic communication and strong partnerships.

We report on the results of these conservation efforts in the Dinaric Mountains and the Southeastern Alps, including the integration of the translocated lynx into the population and local ecosystems, as well as the efficacy of different release methods (soft vs. hard releases). We also compared survival, mortality causes, reproduction and predation among translocated and remnant lynx. In parallel, we conducted systematic camera-trapping and non-invasive genetic monitoring of the population at a transboundary level to continuously measure the population-level impact of translocations on demography and genetic diversity. Results of this monitoring guided adaptive decision-making on the annual basis throughout the translocation process. Furthermore, we used structured questionnaires to measure public and stakeholder support before, during and after the translocation process. This resulted in a comprehensive and holistic translocation effort that transcended jurisdictional boundaries and engaged stakeholders in meaningful and sustainable conservation of a large carnivore in a human-dominated landscape.

## 2. Methods

### 2.1 GPS tracking, translocations, and integration of translocated animals

In total, 50 lynx were equipped with GPS-GSM/Iridium-VHF telemetry collars with timer-controlled drop-off system (Vectronic Aerospace GmbH, Germany and Followit AB, Sweden) in the northern Dinaric Mountains and the Southeastern Alps in Italy, Slovenia, and Croatia (44°–46 ° N, 13° –16° E; Fig. 1). These included all translocated lynx (n=22; 2019-2024), some of their offspring (i.e. first-generation offspring, hereafter “F1”, with at least one of the parents being a translocated lynx; n=10; 2020-2024) and some of the lynx from the remnant population (hereafter “remnant lynx”; n=18; 2006-2024). The GPS fix schedule varied between individuals, seasons and research focus (mean: 3 fixes per day; range: 0.5-96 fixes per day; see Dataset S1 for details). Some of the lynx were re- captured for collar replacement in order to prolong the tracking period. Capturing and immobilisation was done using standard protocols (for details seeKrofel et al. 2013). GPS-tracking periods among individual lynx ranged between 1 and 1,444 days (Dataset S1).

**Fig. 1.**
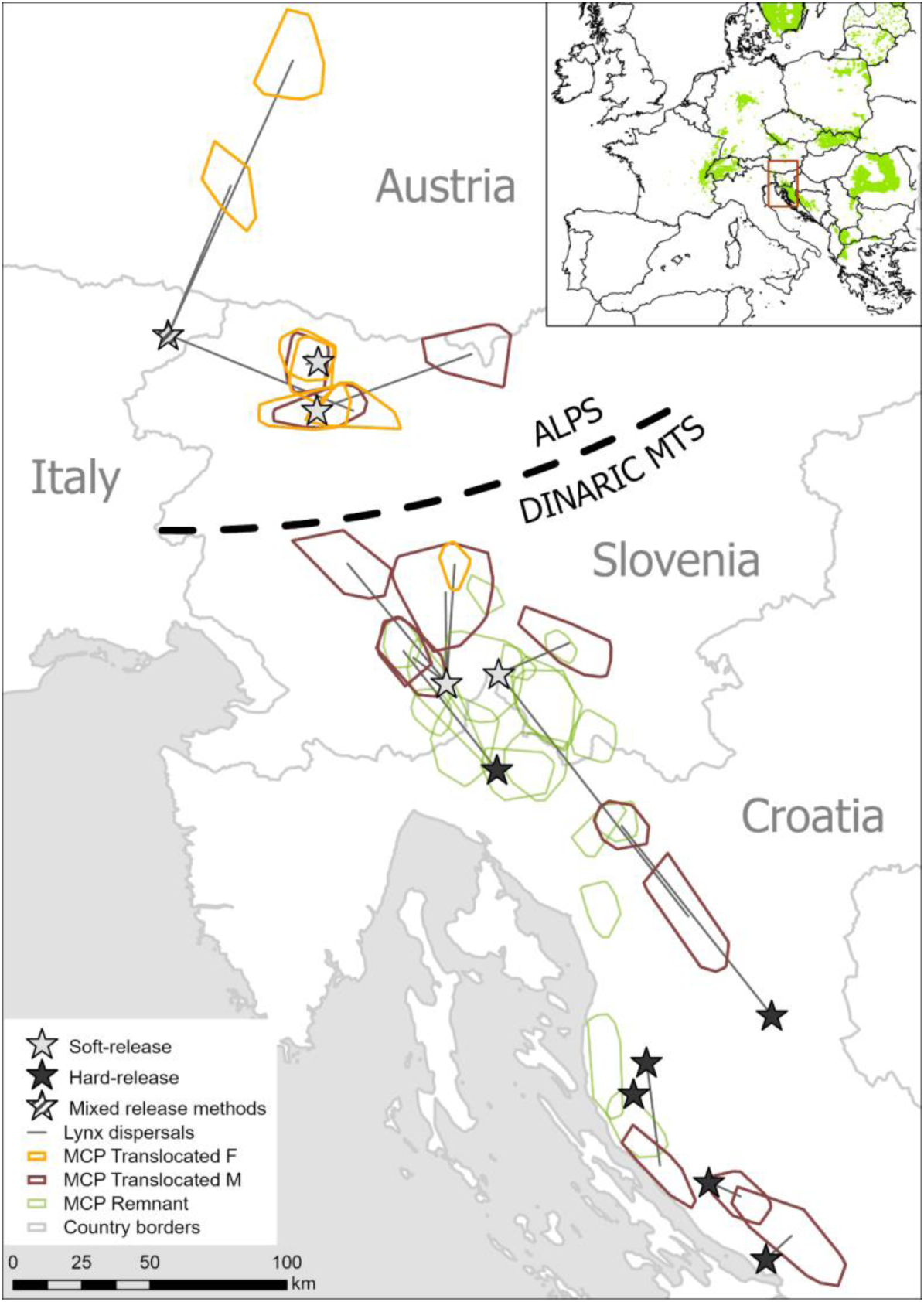
Map depicting the Dinaric and Alpine study areas with the release sites (stars), home ranges of the established translocated lynx (orange = females, brown = males) and GPS-collared remnant lynx (green). Straight black lines connect release sites with home-range centroids of the established translocated lynx that dispersed after release. Note that only a small proportion of the remnant lynx were collared. Not shown on the map are the GPS-tracked F1 lynx and the translocated lynx that did not establish home ranges. The insert map shows location of the study area in Continental Europe with lynx distribution in 2016 (source: www.lcie.org).

All translocated lynx were first quarantined in the country where they were captured (minimum three weeks in Romania and Slovakia and minimum one week in Switzerland). After quarantining, the lynx were transported and hard released (i.e., without an acclimation period) in Croatia and Italy or soft-released with an acclimation period at the release site in Slovenia and Italy (see Supplementary Methods, Section 1.1 and Dataset S1 for details on capturing, quarantine and soft releases). We considered translocated lynx as established when exhibiting a polygonal movement (Topličanec et al. 2022). If they dispersed <5 km (i.e. approximate radius of female home range) from the release site, we considered them as established in the release area. We considered lynx as integrated in the population if they established a permanent home range overlapping with conspecifics of the opposite sex for >1 year and/or were confirmed to have reproduced at least once.

To understand the integration of the translocated lynx into local ecosystems, we focused on two interspecific interactions that were reported as the most important for lynx ecology in Dinaric Mountains, i.e. predation and kleptoparasitism (Krofel 2012). We compared predation parameters (feeding time and inter-kill intervals) of the translocated (N=20) and remnant (n=7) individuals that had suitable GPS fix schedules to predict kill sites following the methodology described in Oliveira et al. (2023). Kills were identified with GPS location cluster (GLC) analysis with cluster criteria of 200-m spatial buffer, two days of spatial window, and a minimum number of two GPS fixes (Oliveira et al. 2023). We visited a subset of the clusters in the field (n=253) to determine prey composition (Krofel et al. 2014) and measured time needed for the released lynx to make the first kill (Topličanec et al. 2022). We deployed camera-traps on fresh kills (n=62) to monitor prey consumption and detect scavengers benefiting from lynx kills (Krofel et al. 2019). Because kleptoparasitism by brown bears (*Ursus arctos*) was observed to play important role for lynx energetic input in the Dinaric mountains, where bear densities are much higher than in the Alps (Krofel and Jerina 2016), we compared bear kleptoparasitism rates (proportion of lynx kills found by bears) and feeding time for kills of translocated lynx in the Dinaric Mountains and the Alps (Krofel et al. 2012).

### 2.2 Survival and mortality

We categorized lynx status at the end of GPS-tracking periods following Andren et al. (2006) as: alive, disappeared, suspected poaching, confirmed poaching, road-kill, and natural mortality (definitions are provided in the Supplementary Methods, Section 1.1). Suspected poaching, confirmed poaching, and road-kills were considered as human-caused mortality. Any potential mortality was immediately investigated when collars emitted a mortality mode or lynx were immobile for several days. All dead lynx that were found were necropsied.

We estimated survivorship based on lynx status at the end of each GPS-tracking period (Dataset S1). We excluded data for five lynx with unknown status (category: disappeared). One remnant lynx was recaptured 10 years after his first capture, and thus we considered the two tracking periods independently. We considered a total of 45 tracking periods to estimate survival rates (Dataset S1). We used the product-limit (i.e., Kaplan-Meier) estimator (Kaplan and Meier 1958) applied using the R package “survival” (Therneau et al. 2021) to determine survival rates for the three lynx groups (Fig. S2).

### 2.3 Reproduction

Reproduction of translocated females was detected by visiting den sites identified through GLC analysis (Krofel et al. 2013) or with camera-trapping (see 2.5). Reproduction of translocated males was confirmed by genetic analyses of parenthood of the sampled kittens (see 2.6) or assumed when presence of a family group (i.e. female with dependent kittens) was detected with camera traps inside established male home ranges.

Litter sizes of translocated and remnant lynx were estimated using camera-trap images of family groups detected between August and April. We used the largest number of kittens observed for a given female and season for further analysis. We used Wilcoxon rank-sum test to compare litter sizes between translocated and remnant lynx.

### 2.4 Population density and abundance

We used camera-trapping and spatial capture-recapture (SCR) analysis to assess changes in the density and abundance of the Dinaric lynx subpopulation in Slovenia and Croatia during the four consecutive years of the reinforcement process (2019-2023), following methodology described in Fležar et al. (2023) (see also Supplementary Methods, Section 1.3 for further details). We classified the camera trapping sites (hereafter ‘location type’), based on their main characteristics; 1) lynx scent-marking sites; 2) forest roads and 3) other sites (Supplementary Methods, Section 1.3 and Fig. S4). We selected at least two camera trapping sites in a potential home range of a studied population (Royle et al. 2011), with a 95% average MCP home-range size of adult female lynx 97 km^2^ (Dataset S1). Although some camera traps were active throughout the calendar year, we limited the camera-trapping data for SCR modeling to a period from August 15th to February 15th (Table S4) to meet the demographic closure assumption. However, to calculate population turnover (Figure S8), we used all available records of independent individual lynx.

Lynx population density, baseline detection rate and spatial scale parameters were estimated with maximum likelihood SCR models (Royle et al. 2011) using R package “oSCR” (Sutherland et al. 2019). We ran multi-session models with four sessions defined as respective survey years in the Dinaric Mountains. We tested the effect of the local behavioral response (‘b’) (Fležar et al. 2023; Royle et al. 2011) and the additive effect of survey year and sex on baseline detection rate and the spatial scale parameter, as suggested by Goldberg et al. (2015). Sex was included as a binary covariate (female as reference category) and survey year as factorial covariate. We included the additive effect of location type as a categorical three-level covariate with marking site as reference category (Table S6), following Fležar et al. (2023).

For each survey year, we defined the extent of the effective sampling area, i.e. the “state space” with the buffer width of 15 km and the resolution of buffer cells 2.5 x 2.5 km (Fig. S4; Fležar et al. 2023; Royle et al. 2011). We ranked candidate models based on Akaike Information Criterion (AIC) and their predictive power (AIC weight). The model with the best fit and highest predictive power was used to calculate the density and abundance of the lynx in the Dinaric mountains (i.e. without the Alpine part) for each survey year.

We deployed camera-traps at 225-329 sites per survey year in 2019-2023, resulting in a total of 116,902 camera-trapping days. For SCR modeling, we used lynx data from 1021 independent occasions to create 234 individual capture histories (Table S5). Model with the most support included survey year, local behavioral response, sex and location type as effects on the baseline detection probability parameter, and sex as an effect on spatial scale parameter (Table S3). Males were detected roughly twice as much as the females, and the camera traps deployed at marking sites had approximately 2-times and 3-times higher probability of detecting lynx than on roads and other locations, respectively (Table S7).

### 2.5 Genetic monitoring and inbreeding assessment

We used a dedicated laboratory for DNA extraction and PCR setup from non-invasive and historic genetic samples. For genotyping, we used a set of 19 microsatellite markers (marker list together with protocols are provided in the Supplementary Methods, Section 1.4). Genetic diversity was measured as the observed heterozygosity (*Ho*) and Nei’s unbiased expected heterozygosity (*He*) (Nei 1978). We used a travelling window analysis (Sindičić et al. 2013) to explore the erosion of genetic diversity caused by genetic drift in the Dinaric population with 40 samples as the width of the window. All analyses were programmed in R, genetic data were handled with the R package ‘adegenet’ (Jombart et al. 2008).

F1 individuals were detected through the private alleles not found in the Dinaric population prior to the translocation (26 private alleles on 11 loci) and they were assigned to possible parents through simple exclusion.

Altogether we analyzed 750 genetic samples from the Dinaric Mountains and the Southeastern Alps collected in the 2010 - 2023 period, including the translocated lynx, and 204 historical genetic samples from 1979-2010 (Supplementary Results, Section 2.3). In total, we identified 238 individuals, and used genotypes of 228 individuals (1979-2010 = 88, 2011 - 2016 = 25, 2017 - 2023 = 117) for downstream analyses. Some animals were excluded because of incomplete field data (N=3), and we excluded the translocated animals that we know had died before reproduction or had no chance to reproduce (n=7).

We used Wright’s hierarchical structuring of inbreeding (Wright 1931) to estimate the dynamics of inbreeding of the Dinaric population relative to the source population from the Slovakian Carpathians. We used the term ‘*effective inbreeding’ (Fe)* (Frankham et al. 2010), where, with *H_Din_* being heterozygosity in the Dinaric lynx and *H_SK_* being heterozygosity in the source population in Slovakia (estimated to He = 0.592, using the same markers and 60 individuals). We estimated the expected inbreeding depression *δ* using 12 diploid lethal equivalents (2B) (O’Grady et al. 2006). Remaining relative fitness compared to Slovak lynx was calculated as 1 − *δ* (for details see Supplementary Results, Section 2.3).

To understand the effect of translocations, we explored three scenarios of inbreeding development after the reinforcement: excluding the effect of translocations (only remnant lynx), including only lynx translocated to Dinaric Mountains and their offspring (isolated stepping stone scenario), and including all translocated animals assuming future merging of the Alpine stepping stone with the Dinaric subpopulation (fully-connected stepping stone scenario).

### 2.6 Public and stakeholder attitude surveys

We measured support for lynx conservation and translocations among the general public and hunters through quantitative public attitude surveys, using the same methodological framework outlined in Majić et al. (2011). Surveys were conducted at the project’s inception (2019), midway point (2021) and conclusion (2023). Random samples of adult (>18 year old) residents of the project area were obtained either through panel samples and implemented online (Croatia, Italy) or through sampling from the register of residents and implemented via regular mail (Slovenia, Croatia), while hunters were sampled by mail via hunting organizations and personally at events (see Supplementary Methods, Section 1.5 for details).

## 3. Results

### 3.1 Translocations, post-release movement and integration

A total of 22 lynx (7F, 15M) were translocated from Romania (2F, 10M), Slovakia (3F, 5M) and Switzerland (2F). These lynx were released to reinforce the remnant lynx population in the Dinaric mountains in Slovenia (1F, 5M) and Croatia (6M), and to create a new stepping-stone subpopulation in the Southeastern Alps in Slovenia (3F, 3M) and Italy (3F, 1M) (Fig. 1, Dataset S1). All releases were supported and partly conducted by hunters from local hunting organizations or protected area managers, all of whom were directly involved in all phases of the translocation process (Supplementary Methods, Section 1.5.1).

Of the 22 translocated lynx, we considered 15 (68%) to have been successfully integrated into the population, 6 (27%) died or disappeared before reproducing, and the status of the remaining one animal is unknown (still in dispersal phase) (Dataset S1). Among the soft-released (n=14) and hard- released animals (n=8) (Fig. S1), integration was successful for 71% and 63% of the lynx, respectively. Soft-releases of lynx in the Slovenian Dinaric Mountains and Alps had the greatest population integration success (83% in both areas; Dataset S1). Overall, 32% of the translocated lynx established permanent home ranges in the release area (<5 km from the release site), including 43% of the soft- released and 13% of the hard-released lynx. We observed the lowest dispersal rates of lynx released in the Slovenian Alps, where 83% established permanent home ranges in the release area (Fig. 1, Dataset S1). One of the males released in the Slovenian Dinaric Mountains moved to the Alps and back, indicating opportunity for functional connectivity between the two study areas. The median (± IQR) distance between the release site and centroid of established home ranges was 23.20 ± 34.97 km (n=19; mean = 27.91 km). The median distances were 41% shorter for the soft-released lynx (19.57 ± 37.58 km; n= 13) than for the hard-released lynx (32.92 ± 33.96 km; n=6), although the difference was not statistically significant (Kruskal-Wallis χ^2^ = 0.77, p = 0.38).

On average (± SD) translocated lynx made the first large kill 8.75 ± 6.85 days after release (n = 12; Dataset S1). The mean inter-kill intervals (± SE) for large prey of the translocated lynx (4.37 ± 0.12 days; n=621 predicted kills) were almost identical to those from the remnant population (4.38 ± 0.14 days; n=385; Kruskal-Wallis χ^2^, p=0.88) (Dataset S1). Translocated lynx mainly (83%) killed roe deer (*Capreolus capreolus*) and we observed 19 species of scavengers feeding on their kills (see Tables S1 and S2 for full lists of prey and scavenger species). Brown bear kleptoparasitism rates were much lower in the Alps (4.0%) compared to the Dinaric mountains (32.3%). Accordingly, we observed 15% longer feeding times of the translocated lynx in the Alps (mean ± SD: 2.41 ± 0.08 days) compared to the Dinaric Mountains (2.09 ± 0.07; Kruskal-Wallis χ^2^, p = 0.006).

### 3.2. Survival and mortality causes

Among the 50 GPS-tracked lynx, 33 survived the entire GPS-tracking tracking periods, five disappeared and 11 were confirmed or suspected to die. Survival was the highest for the offspring of translocated lynx (F1; n=10), followed by translocated individuals (n=22) and lowest for remnant lynx (n=18), although the differences in the survival probability were not significant (Kaplan-Meier, p=0.52; Fig S2). Persistence seemed particularly low for the four lynx that dispersed into Austria or along the Austrian border (Fig. 2). Confirmed or suspected mortality was the highest for the remnant lynx (33%), followed by the translocated (18%) and F1 lynx (11%) (Dataset S1). All confirmed or suspected mortalities among the translocated and F1 lynx (n=5) appear to be human-caused (confirmed or suspected poaching). Among the mortalities recorded among the remnant lynx (n=6), 33% were probably human-caused (road-kill and suspected poaching) and 67% were natural. Among natural mortalities we recorded two cases of heart failure connected with congenital atrial septal defects, one case of pneumonia and in one case the exact cause was unclear. Moreover, we detected additional morphological deformations or their manifestations possibly linked to inbreeding among the remnant lynx (heart murmurs, skeletal deformations and double ear-tufts on the same ear instead of normal, single ear-tuft; Fig. S3). No deformations or natural mortalities were observed among the translocated or F1 lynx.

**Fig. 2.**
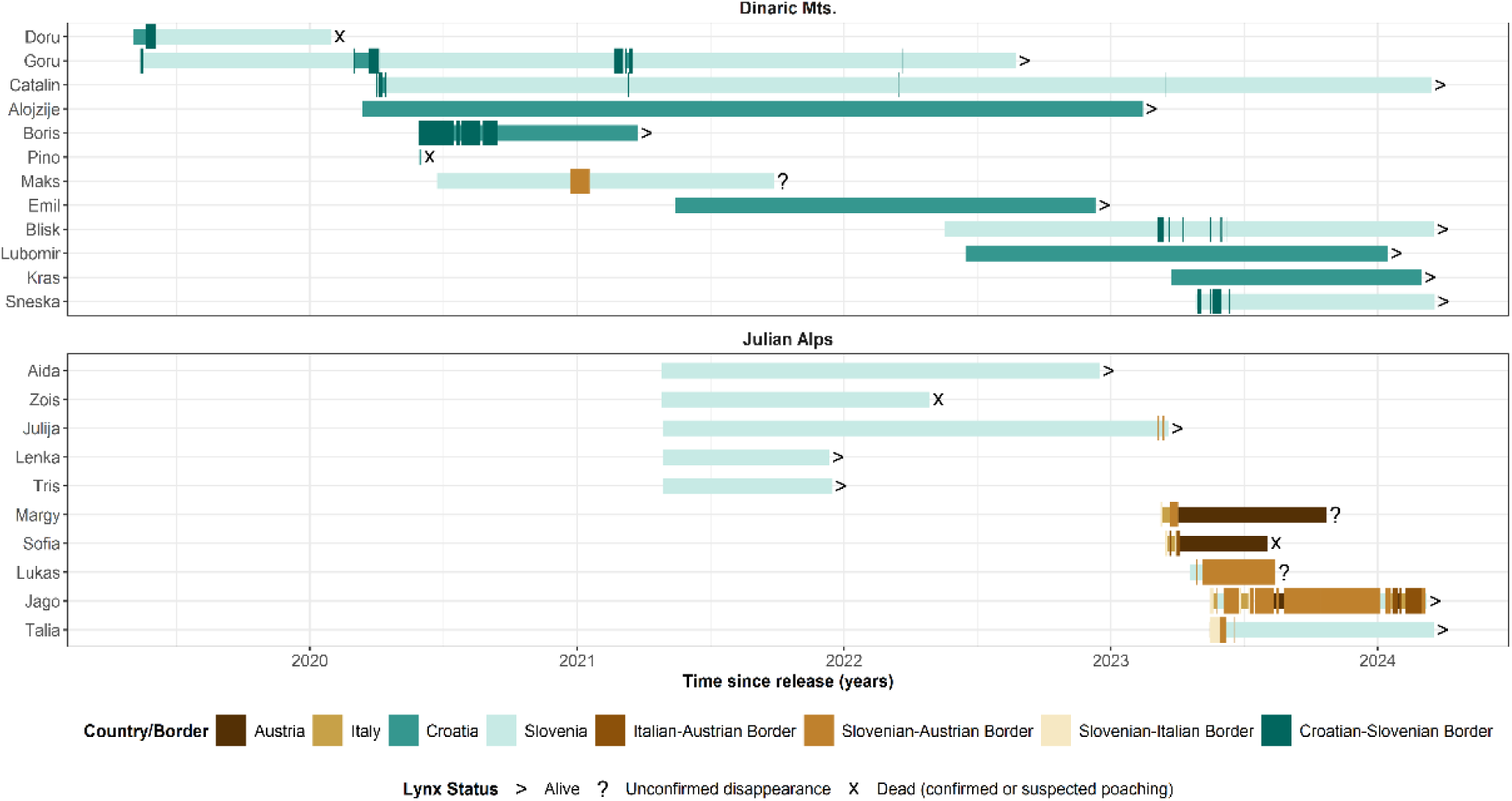
Spatio-temporal representation of lynx movements within GPS-tracking periods and status at the end of tracking periods for 22 lynx translocated to the Dinaric Mountains (above) and the Julian Alps (below). Colors and width of lines indicate location at given countries or border zones (i.e. at 5-km buffer around the border).

**Fig. 3.**
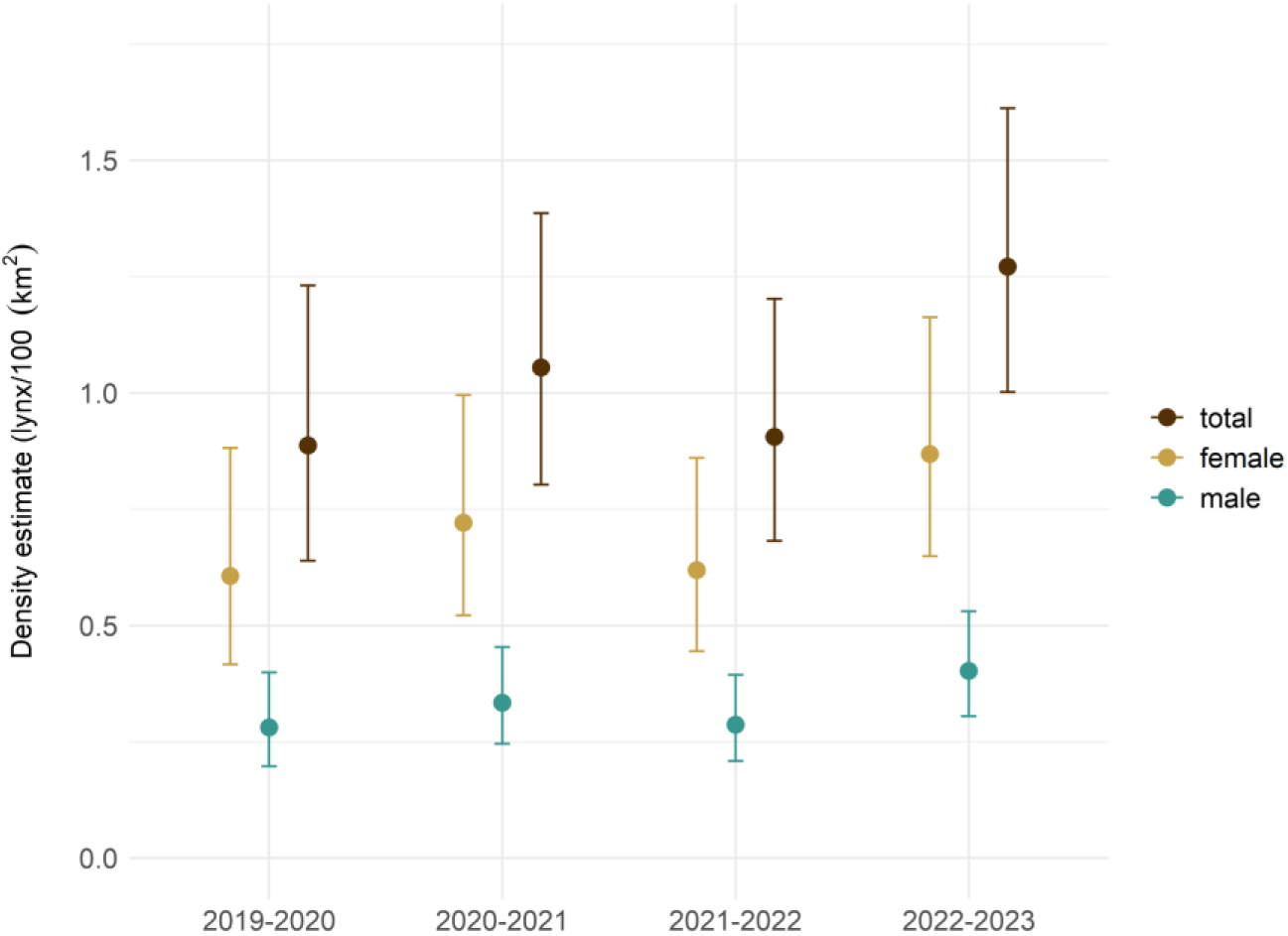
Changes in density estimates for the lynx in the Dinaric Mountains during the reinforcement process (2019-2023). Estimates (with 95% confidence intervals) are shown for each survey year for females, males and all independent lynx.

**Fig. 4.**
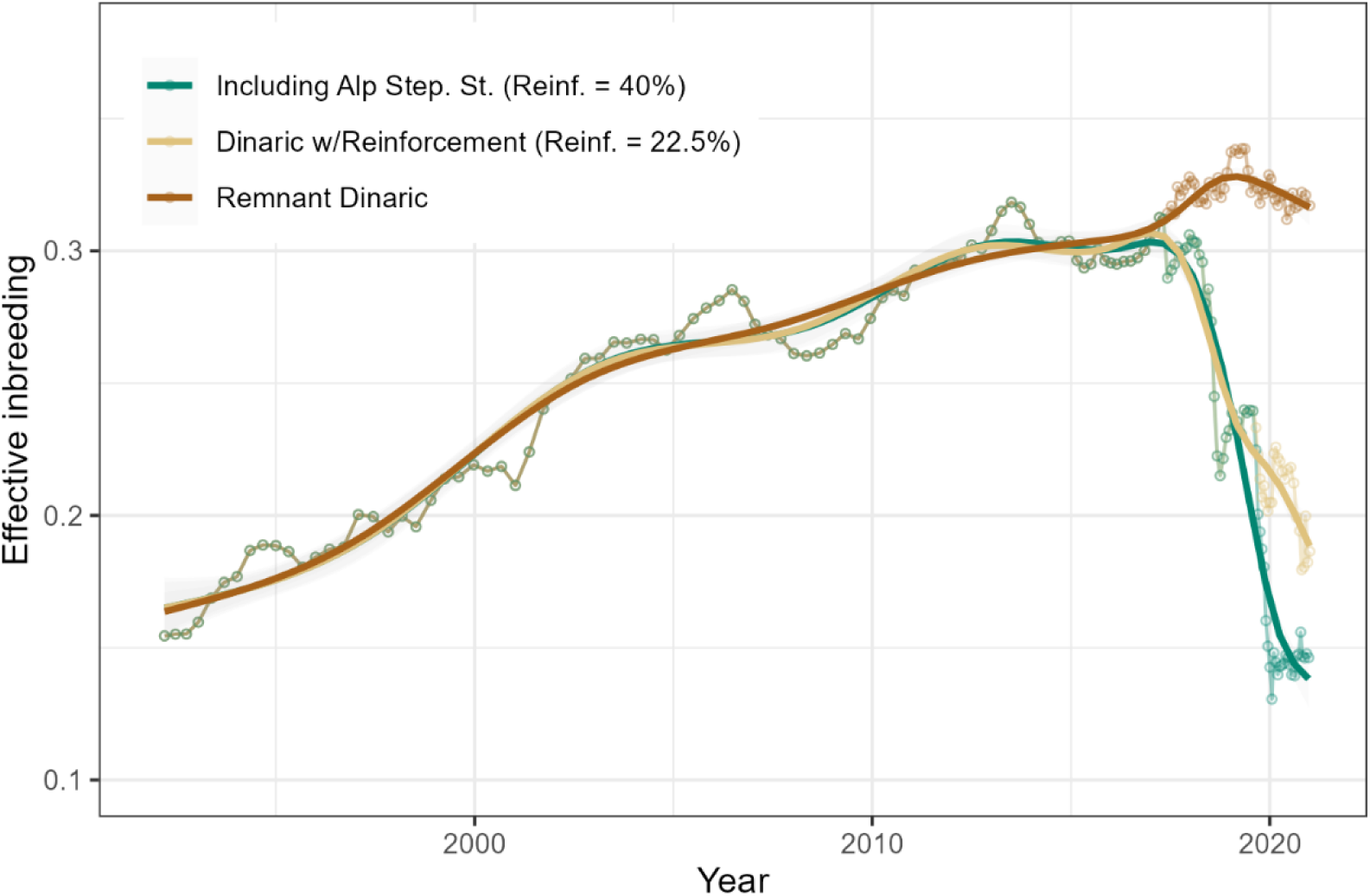
Effective inbreeding (Fe) of Dinaric lynx relative to the population in the Slovak Carpathians (source for the 1973 reintroduction), calculated using a 40-sample traveling window. Remnant Dinaric: calculated without the translocated lynx and their offspring, exploring situation without the effect of translocations; Dinaric w/Reinforcement: including lynx translocated to Dinaric Mountains and their offspring; Including Alp Step. St.: with all translocated lynx, including the Alpine stepping stone subpopulation. The “Reinf.” value indicates the proportion of translocated animals and their offspring in the final traveling window (right end of the graph).

### 3.3 Reproduction

Among the 22 translocated lynx, 13 (59%) reproduced by 2024 and three more individuals have the potential for future reproduction (Dataset S1). We were able to confirm the parentage of translocated lynx in 10 out of 23 litters detected by camera trapping. We recorded a total of 54 kittens with presumed or confirmed parentage from translocated lynx. This is likely an underestimate, because males may have sired additional offspring during extra-territorial mating excursions. Such excursions were recorded in 10 out of the 15 mating seasons of the nine GPS-tracked translocated males. Also not included in the reported number of kittens is reproduction of F1 lynx (one litter confirmed so far).

In total, we obtained data from 97 lynx litters by camera-trapping. When at least one of the parents was presumed or confirmed to be a translocated or F1 lynx, litter sizes were 37% larger compared to the litters with both parents from the remnant population (F1 litters: 2.21 ± 0.78 kittens, n=24; only confirmed F1 litters: 2.30 ± 0.95 kittens, n=10; remnant litters: 1.62 ± 0.68 kittens, n=73; p<0.001).

### 3.4 Changes in the population density and abundance

During the 4-year reinforcement process in the Dinaric Mountains, the mean lynx population density increased by 44.3% (from 0.88 [95% CI: 0.63-1.23] independent lynx / 100 km^2^ in 2019-2020 to 1.27 [1.00-1.61] in 2022-2023), with the highest increase in the last survey year (Fig. 2, Table S3). This increase occurred without any major changes in the effective area surveyed, which ranged between 12,206-12,350 km^2^ per survey year (Fig. S4, Table S5). The estimated densities correspond to a 42% increase in abundance from 110 (95% CI: 79-152) adult lynx in 2019-2020 to 156 (123-198) adult lynx in 2022-2023 in the state space (Table S5).

Despite the population growth, we observed a high turnover rate in the population, which remained similar throughout the survey period (34-36% of individually-identified lynx were detected in the following year; Fig. S8).

### 3.5 Impact on genetic diversity and inbreeding

Without the translocated lynx, inbreeding would reach 0.32 at the end of the project, with expected inbreeding depression of δ = 0.85 (i.e. the fitness of Dinaric remnant lynx is expected to be 15% of those in the source population). It is difficult to precisely estimate the effect of the reinforcement as it will take several generations for allelic frequencies to stabilize and the population to get into the Hardy – Weinberg equilibrium (Cornuet and Luikart 1996), but we can already see that this effect is considerable. Even assuming no connectivity with the Alpine stepping stone subpopulation, inbreeding in the Dinaric lynx would be around 0.19 and inbreeding depression δ = 0.68 at the end of the translocation process, suggesting that fitness already more than doubled due to the reinforcement. If we assume a full connectivity with the Alpine stepping-stone subpopulation, inbreeding would drop to 0.08 when translocated animals and their offspring form around 40% of the population, with expected inbreeding depression dropping to δ = 0.39.

### 3.6 Maintaining public and hunter support

Public and hunter support for lynx conservation remained high throughout the translocation process. Respondents’ commitment to lynx conservation, as indicated by their bequest value orientation (i.e. value that individuals place on the ability to pass a resource to the future generations), remained high among general public and hunters until the final year (2023), with the highest levels of support observed among the Croatian and Slovenian respondents (Fig. 5).

**Fig. 5.**
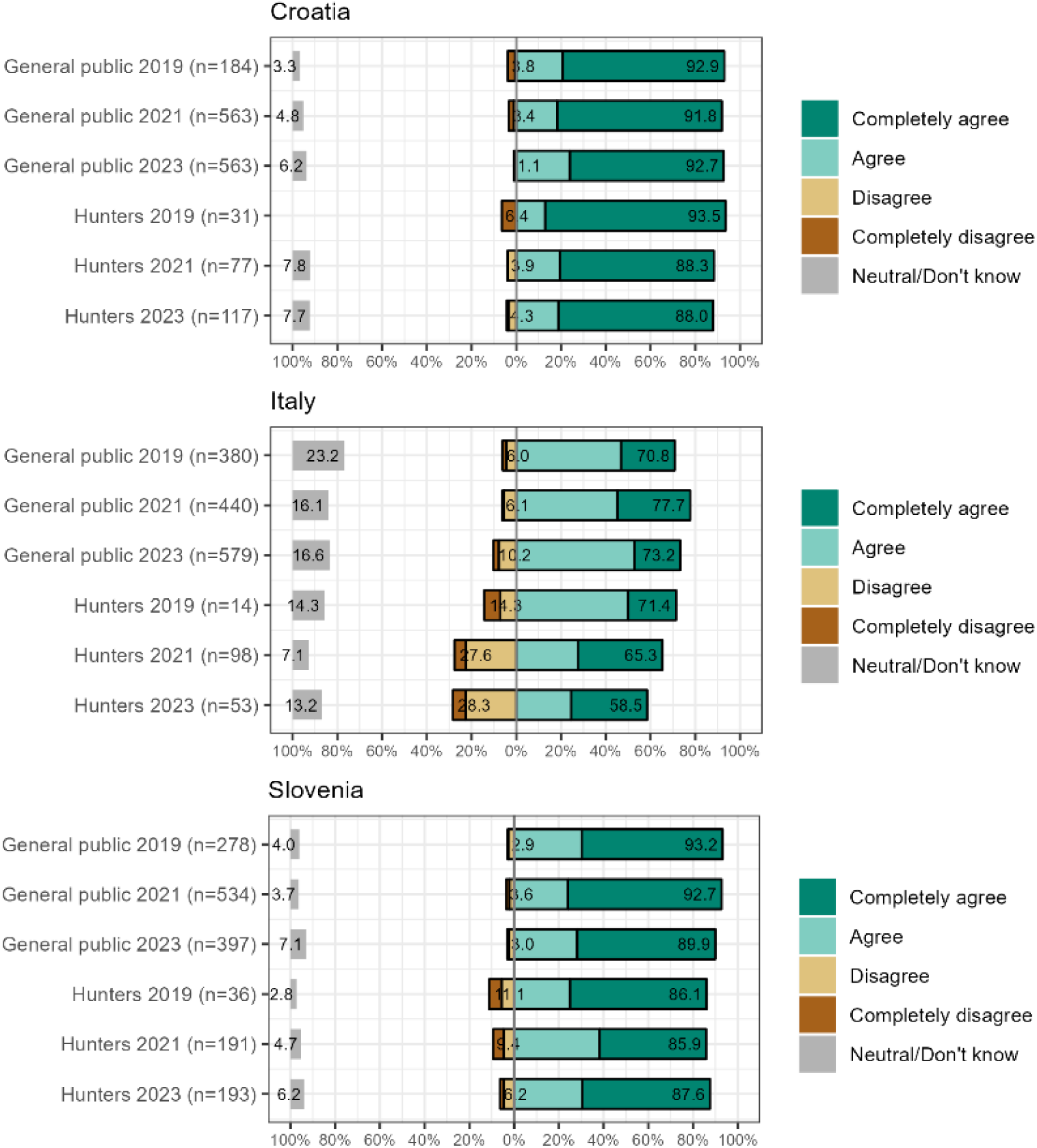
Responses to the statement: “It is important to maintain lynx in Croatia / Italy / Slovenia for the future generations”. Respondents answered for their respective countries.

Additionally, support for translocations was initially high and increased during the mid-project period. As the number of lynx released increased towards the end of the project, there was a slight decline in support among the Italian and Croatian respondents, although the majority still expressed support for the translocation efforts (Fig. S10 and Dataset S2).

## 4. Discussion

Rescue of the Dinaric lynx population demonstrates the feasibility of using translocations to recover a highly-inbred large carnivore in a human-dominated landscape and across national borders.

Besides carefully designed translocations, the efforts to communicate regularly and transparently with the public and engage key stakeholders in the project implementation helped make the effort a success. Our study also provides a rare example of a comprehensive monitoring program implemented on a transboundary level during a translocation effort. We monitored both biological and social parameters before, during and at the end of translocation process, thus ensuring that we could evaluate and adapt management options throughout the process and ensure a holistic understanding of the implemented reinforcement and reintroduction. We suggest that this project may serve as a blueprint for planning future efforts to re-establish healthy populations of large carnivores in human-dominated landscapes.

### 4.1 Mortality, reproduction and integration success

Lynx that were soft-released showed higher success rates of integration into the remnant population and shorter post-release movements than hard-released lynx. This is in line with previous research on translocations of carnivores and other terrestrial animals (Resende et al. 2021; Thomas et al. 2023). However, it also appeared that the presence of conspecifics of the opposite and same sex played an important role for post-release behavior (e.g. comparison between soft-released animals in the Slovenian Alps and Dinaric Mountains), which warrants further analysis.

Natural causes were dominant among the remnant animals that were found dead and especially congenital heart defects with other morphological deformations indicate possible effects of the inbreeding depression. Mortality was lower among translocated lynx, where the main cause was poaching. Especially important concern are rapid disappearances of lynx that moved across or to the border with Austria, which aligns with high poaching rates of lynx and other large carnivores reported for this region (Heurich et al. 2018; Kaczensky et al. 2011; Molinari et al. 2021). To address the threat of poaching, LIFE Lynx project invested considerable efforts in partnership with hunters (for details see Supplementary Methods, Section 1.5), including establishing the first specialized anti-poaching police team in the region.

Six months after releases, survivorship of translocated lynx (86%) was higher compared to average survivorship (66%) reported by Thomas et al. (2023) for recent large carnivore translocations. The majority of translocated lynx also succeeded in reproducing and we observed significantly higher reproductive success compared to the remnant lynx, probably related to lower inbreeding levels. An important additional factor that likely contributed to high rates of survivorship was that we used only wild-born individuals who had innate hunting ability. This is supported by no signs of reduced kill rates and the relatively short periods needed to capture the first large prey. This prevented starvation, which is often a leading cause of mortality among translocated predators (Devineau et al. 2010). Bear kleptoparasitism rates for translocated lynx in the Dinaric Mountains were similar to the remnant population (Krofel and Jerina 2016), but they were considerably lower among the lynx translocated to the Alps. This could explain longer feeding times (and therefore higher food intake) in the Alps, which could from this perspective represent a more advantageous lynx habitat. Overall, the observed interspecific interactions involving translocated animals were similar to the remnant lynx (Krofel et al. 2012, 2014, 2019), which suggest that most of the translocated lynx became quickly integrated in the local ecosystems.

### 4.2 Impact on the population demography

After the 2000s, the population experienced a drastic decline and became functionally extinct in the Alpine study area (Fležar et al. 2021). After the reinforcement was initiated in 2019, the population trend in the Dinaric Mountains reversed and the lynx abundance increased by more than 40% in the following three years. The increase was mainly connected with increasing density and was non-linear in time with a major change detected in the last survey year. This time lag was likely due to gradual integration of new animals in the population and time needed for their offspring (>50 kittens detected) to mature to independent animals, which are considered in the camera-trapping monitoring. Current population densities already surpass many of the density estimates from other reintroduced lynx populations in Europe and approach those of the source population (reviewed in Fležar et al. 2023). With saturation of the core area, we expect that in the following years the population will start expanding to the adjacent areas with suitable habitat.

Measuring a large-scale demographic impact of translocations is still rare (Taylor et al. 2017), mainly due to considerable logistic constraints and effort required to conduct transboundary monitoring over vast ranges occupied by large carnivore populations (Fležar et al. 2023; Tourani 2022). Our study demonstrates that citizen science involving close partnership with local amateur and professional hunting organizations and protected area managers can be an efficient approach for obtaining multi-year datasets for a robust population density assessment of an elusive large carnivore. This approach requires considerable effort in communication and coordination, but at the same time it results in increased trust in monitoring results and increased support for conservation measures among the involved stakeholders. Furthermore, in combination with intensive international collaboration, such camera-trapping surveys can become an achievable goal even over large (>10,000 km^2^) scales and spanning across multiple countries and management jurisdictions, which can be major limitation for carnivore translocations (Port et al. 2024).

A priority for the future is to expand this monitoring to Bosnia and Herzegovina, where very limited information about lynx is currently available. Besides, this country is essential for establishing connection with the critically endangered and probably inbred Balkan lynx population towards the south-east (Melovski et al. 2022).

### 4.3 Impact on genetic diversity and inbreeding

The genetic erosion of the Dinaric lynx population since the 1973 reintroduction and critical levels of inbreeding (Fe > 0.3) resulted in a severe inbreeding depression (expected δ = 0.85), suggested also by the observed congenital heart defects and morphological deformations. The post-reinforcement situation shows a dramatic improvement, with a considerably increased expected heterozygosity, and the related expected fitness more than doubling since the reinforcement. This is corroborated by other field data showing encouraging signs of recovery (increased population size, larger litter sizes, and decreased natural mortality). We can expect that inbreeding in the population will continue dropping as offspring of the translocated animals spread through the population and allelic frequencies stabilize (Cornuet and Luikart 1996).

Over the next decade, the status of the population will depend on the continued reproductive performance of the translocated animals and their offspring. In the well-studied case of the reinforcement of the Florida panther (*Puma concolor coryi*) population, considerable heterosis (fitness advantage of outbred animals) was observed in the F1 animals, which contributed to rapid expansion of the introduced genes in the population (Johnson et al. 2010). After the reinforcement, panther numbers increased threefold, heterozygosity doubled, survival and fitness measures improved, and inbreeding correlates declined significantly. We already started observing similar changes in the Dinaric lynx. Moreover, as most of the translocated animals originated from another part of the Carpathian lynx range (Romania) than the remnant population (Slovakia), the resulting F1 offspring are even more outbred than the lynx in the source populations. A continuous monitoring program should be a priority to keep track of future developments, especially since the population still remains small, isolated, and bottlenecked. In the long term, this will require creating a large Central European metapopulation or conducting further periodic translocation of outbred lynx.

### 4.4 Maintaining public support

Any predator translocation or increase in carnivore populations can result in intensified public concerns (Treves and Karanth 2003; Wilson 2018), thus gradual erosion of public support and opposition of key stakeholders (hunters) were among the main concerns in the planning phase of lynx translocation. Various stakeholder engagement efforts along with intensive communication campaigns within the LIFE Lynx project likely contributed to sustaining high support for lynx conservation among the local hunters and general public throughout the translocation process and population increase. Although the majority was also in favor of the translocations throughout the process, this support has slightly tapered off towards the end of the project in Italy and Croatia, possibly reflecting the lower perceived need for further translocations, when population was no longer in the immediate risk of extinction. With the lynx population steadily recovering, future stakeholder engagement strategies should pivot towards (1) addressing stakeholder-specific and country-specific concerns regarding the implications of the lynx recovery, and (2) effectively communicating the necessity of long-term measures to ensure the establishment of a viable lynx population at a transnational level.

### 4.5 Future perspectives

Central Europe is a highly-fragmented landscape and none of the existing forest complexes are large enough to host a viable isolated lynx population. Therefore, the establishment of a functional transboundary metapopulation is needed to mitigate the negative effects of habitat fragmentation and ensure long-term viability (Mueller et al. 2022). Genetic restoration of the Dinaric population and the creation of a new stepping-stone subpopulation in the Julian Alps reported in this study represent a major advance towards reaching this long-term vision. Next step is the creation of additional stepping stones that could eventually re-establish connectivity across the Alpine arc with (Western) Alpine and Jura populations in Switzerland and France (Molinari et al. 2021). However, the high poaching rates in Austria and neighboring regions in the Alps will likely be a major obstacle in achieving this goal. This calls for a targeted, multi-sectoral approach, which considers the practice of stakeholder engagement reported here and acknowledges the given cultural context.

As UN decade on ecosystem restoration progresses, numerous carnivore restoration projects are currently being planned or already initiated globally, including efforts to return lynx to the Great Britain and several parts of Germany, gray wolves to Colorado, USA, cheetahs to India, and leopards to Saudi Arabia. Lessons learned from our comprehensive conservation endeavors coupled with the multidisciplinary monitoring of biological and social factors could provide an important example to facilitate these carnivore restoration efforts.

## Acknowledgements

We are grateful to the LIFE Lynx Project personnel, as well as hundreds of hunters, foresters, veterinarians, technicians, students and volunteers for their invaluable help with the fieldwork and data collection. This study was co-funded by the European Commission under the LIFE Programme (LIFE16 NAT/SI/000634 LIFE Lynx project) and the Slovenian Research and Innovation Agency (grants J1–50013, P4-0059, and P1-0184). Lynx releases in Italy were financed by the WWF.

