## Supplementary material for "Saving the Dinaric lynx: multidisciplinary monitoring and stakeholder engagement support large carnivore restoration in human-dominated landscape"

**Content**


| | |
|---|---|
| Supplementary Methods | pages 2 – 6 |
| Supplementary Results | pages 7 – 8 |
| Supplementary Figures S1-S10 | pages 9 – 18 |
| Supplementary Tables S1-S8 | pages 18 – 26 |
| Supplementary Datasets S1-S2 | page 27 |
| Supplementary References | pages 28 – 30 |

### 1. Supplementary methods

#### 1.1 Surveys in source populations, translocations, movements, survival and mortality

Before capturing lynx for translocation, source populations were surveyed to confirm that removal of animals needed for translocations would be sustainable and would not jeopardize these populations (for detials see Kubala et al. 2018; Sin et al. 2021).

Translocated lynx were quarantined for a minimum of three weeks in Romania and Slovakia and for a minimum of one week in Switzerland. Protocols used for capturing, quarantine, transport and releases of lynx are available on the project webpage (https://www.lifelynx.eu/protocols).

Soft-release followed an acclimation period at the release site lasting on average 19.1 ± 14.7 days (range: 3-48 days) in Slovenia and Italy (for details see Dataset S1).

The process of integration of each translocated lynx was continuously monitored, which guided us in decisions on where and when to release the next animals. Relatedness with already released animals was also considered in the release decision-making.

Definitions of categories describing lynx status at the end of GPS-tracking periods:
- alive: lynx still tracked at the end of the study or moving when the collar drop-off was triggered
- disappeared: collar signal lost (i.e. no more GSM/Iridium data transmission and no VHF signal found in the lynx home range) before the expected battery lifetime, but no systematic monitoring with camera-trapping or snow-tracking was conducted in the area of disappearance to provide further information; mostly likely because the collar failed or lynx was poached, also possible that lynx died of natural causes in a place where GSM/Iridium and VHF collar signals could not be obtained
- suspected poaching: collar signal lost well before the expected battery lifetime and lynx disappeared from camera-trapping records at about the same time
- confirmed poaching: evidence of poaching found, e.g. shot lynx or destroyed collar
- road-kill: lynx killed in vehicle collision
- natural mortality: dead lynx found without signs of human-caused mortality

Because most of the tracked lynx were adults at the time of the collaring/release (i.e., ≥2 years; 76%, n=36), and the collaring/release took place mostly during winter/spring (96%, n=43), we did not include age and season as covariates in our model of lynx survival. GPS-tracking periods that ended without mortality were right-censored, i.e., the survival time was not observed because the tracking ended before the animal died (collar removed, drop-off, or failure).

#### 1.2 Predation and kleptoparasitism

For each kill, we calculated the feeding time and the search time. We defined the feeding time as the period between when the prey was killed and when lynx abandoned the kill (Krofel et al. 2012). We used feeding time as a proxy of lynx food intake per kill. The search time was defined as the period between the abandonment of current prey item and the next kill. The inter-kill interval (analogous to kill rates) considers the time passed between the start of a given kill and the start of a previous kill, and includes the feeding time (capturing and consuming prey) and searching time (the time spent searching for prey) (Oliveira et al. 2023).

#### 1.3 Camera trapping

##### 1.3.1 Fieldwork

We followed the camera trapping guidelines (Stergar and Slijepčević 2017) for camera set up at three types of locations: lynx scent-marking sites (Allen et al. 2017), forest roads (i.e. unpaved forest roads and logging trails), and other sites (i.e. off-road sites where no indication of lynx scent marking was observed). Camera traps were set up exclusively in forested areas, which is the optimal habitat of lynx in the Dinaric Mountains (Fležar et al. 2023). We used cameras with white flash (CuddeBack X-Change Color Model 1279, Cuddeback, Green Bay, Wisconsin; Reconyx HyperFire 2 Professional White Flash Camera HP2W, Reconyx, Inc., Holmen) or cameras with black (940 nm light wave) or regular infrared light (850 nm light wave; CuddeBack X-Change Color Model 1279, Cuddeback, Green Bay, Wisconsin; StealthCam STC-G42NG, Stealth Cam, Irving; Moultrie M40-i, PRADCO Outdoor brands, Birmingham; LTL Acorn models Ltl-6310WMG and Ltl-6511WMC; Browning Spec Ops Elite HP4, Browning Trail Cameras, Birmingham, Alabama).

In Slovenia, camera traps were set at all three types of locations, while camera traps were primarily set at marking sites in the Northern part of Croatia (Gorski Kotar), and at roads in the Southern part of Croatia (Velebit). We limited the camera trapping data for this survey to the sites located South of the A1 highway in Slovenia. The highway represents a serious barrier to the lynx home range, i.e. telemetry tracking showed no lynx established a territory overlapping the highway, although some dispersing lynx managed to cross it (Krofel et al. 2006, 2021). The data from the Alpine part was not considered for the SCR modeling, as the stepping stone subpopulation was not yet connected to the Dinaric population and only consisted of the known translocated lynx and their offspring at the time of this study. The mean trap spacing (Table S4) and spatial scale parameter calculated by the models (see Results) confirmed that designs in all regions and the entire study area fit the standard recommendation of 2σ (Dupont et al. 2021; Efford and Fewster 2013).

1.3.2 Data preparation

Trained observers (n=9) annotated lynx records with sex, age (juvenile / independent) and individual identity, whenever possible, following the identification guidelines (Choo et al. 2020). Only data about individually-identified independent lynx originating from high-quality records where the coat pattern was clearly visible were used to build individual capture histories. International collaboration between Slovenia and Croatia allowed for uninterrupted exchange of data, including identifying lynx with cross-border territories. This was done also with the help of artificial-intelligence-based tools (www.whiskerbook.org). Records of lynx where identification was not possible were discarded (less than 20% of records each survey year), as well as records of juveniles (Andrén et al. 2006; Duľa et al. 2021). Lynx records and trap deployment data were exported directly from the Camelot software (Hendry and Mann 2017), which was used for camera trapping data annotation, and reorganized to fit SCR analysis.

1.3.3 SCR modelling

Lynx density, baseline detection rate and spatial scale parameter were estimated with maximum likelihood spatial capture-recapture models (Royle et al. 2013) using oSCR package (Sutherland et al. 2019) in R software v. 4.1.0. We ran multi-session models with four sessions defined as respective survey years (2019-2020, 2020-2021, 2021-2022, 2022-2023) in the Dinaric Mountains (i.e. not including the Alpine part). At least 20 spatial recaptures were confirmed for each survey year and for the entire transboundary study area (Dinaric Mountains; Table S5) (Efford et al. 2004).

For each survey year, we defined the extent of the effective sampling area, i.e. the “state space” with the buffer width of 15 km and the resolution of buffer cells 2.5 x 2.5 km, following the recommendations of Royle et al. (2013) and our own findings (Fležar et al. 2023). We restricted the state space following the same criteria as described in Fležar et al. (2023; see also Fig. S5). We ranked candidate models based on Akaike Information Criterion (AIC) with models having ΔAIC ≤ 2 considered having substantial support (Burnham and Anderson 2004), and their predictive power (AIC weight; Johnson and Omland 2004). Among the highest-ranking models, the one with the best fit (lowest AIC value) and the highest predictive power (highest AIC weight) was used to calculate the density and abundance of lynx in the Dinaric Mountains for each survey year.

##### 1.4 Genetics

Tissue samples from dead lynx were stored in 95% non-denatured ethanol and stored at -20°C. Blood was preserved on blood stain cards (Qiagen). These high-quality DNA samples were processed using a manual DNA extraction kit (Sigma GenElute Mammalian Genomic DNA Miniprep Kit) following the manufacturer's protocol.

Non-invasive genetic samples: Urine samples (collected in snow) were stored in DETs buffer, and hair samples were stored in sealed bags with desiccant (silica). Saliva samples were collected with forensic swabs that already have desiccant in the swab tube. Scats were collected in ethanol. Sampling material was prepared and distributed at the beginning of the project, instructions for collecting were presented in dedicated guidelines (Skrbinšek 2017). DNA in non-invasive, historic genetic and eDNA samples is of very low quality and quantity, and contamination (especially with PCR products) is a serious issue. We used a dedicated laboratory for samples storage, DNA extraction and PCR setup. For non-invasive samples we used MagMAX DNA Multi-sample Kit (Thermo Fisher Scientific). The extraction protocol is implemented on a liquid handling robot (Hamilton Starlet), samples IDs are read and handled through barcodes.

We used ten microsatellite markers for individual ID run in a single multiplex: Fca132, Fca201, Fca247, Fca293, Fca391, Fca424, Fca567, Fca650, Fca723, Fca82. The best (reference) sample of each detected animal was amplified using 9 additional markers (F115, F53, Fca001, Fca132, Fca161, Fca369, Fca559, Fca742, HDZ700 (Menotti-Raymond et al. 1999; Menotti-Raymond et al. 2005; Williamson et al. 2002), bringing the total number of studied microsatellites to 19. SRY locus was used to determine sex of the animal. Microsatellites were amplified in 3 multiplexes, using Platinum multiplex PCR Master Mix (ABI). Protocols from (Polanc et al. 2012) were adapted according to the Platinum kit user guide. Good quality tissue and blood samples were re-amplified twice. For non-invasive samples, we used a modified multiple-tube approach (Adams and Waits 2007; Taberlet et al. 1996) with up to 8 re-amplifications of each sample according to the sample's quality and matching with other samples. In the first screening process, each sample was amplified with the 10-marker panel (multiB panel) protocol twice and analyzed on an automatic sequencer (Applied Biosystem ABI 3500 Genetic Analyzer). Results were interpreted using GeneMapper v.6.0. software (Applied Biosystems, USA). Samples that provided no specific PCR products at that stage were discarded. Consensus genotypes were determined using an Access database application programmed by T. Skrbinšek (MisBase, unpublished). Total genotyping success was 47%.

We used Wright's hierarchical structuring of inbreeding (Wright 1931) to estimate the total inbreeding, $F_{it}$. As suggested by (Keller and Waller 2002), in a randomly breeding population ($F_{is} = 0$) the actual inbreeding that would cause inbreeding depression (probability of alleles at a locus being identical by descent) would equal $F_{st}$ between the studied population and metapopulation / entire species. In the case of Dinaric lynx, because the population has been reintroduced from Slovakian Carpathians, the drift component of inbreeding ($F_{st}$) directly indicates the inbreeding of the Dinaric population relative to the source population in the Slovakian Carpathians. We used the term '*effective inbreeding (Fe)*' (Frankham et al. 2002), where $Fe = 1 - H_{Din}/H_{SK}$, with $H_{Din}$ being heterozygosity in the Dinaric lynx and $H_{SK}$ being heterozygosity in the source population in Slovakia. We estimated expected heterozygosity of lynx in Slovakia at $He$ = 0.592, using the same markers and 60 individuals (including the translocated animals). We used this reference to estimate the dynamics of inbreeding in the Dinaric lynx relative to the original source population in Slovakian Carpathians. As the Alpine stepping-stone subpopulation is currently in its first generation and hence completely outbred, it makes no sense to estimate inbreeding for that subpopulation.

A meta-analysis of inbreeding depression in the wild indicated on average 12 diploid lethal equivalents (2B) in wildlife populations (O'Grady et al. 2006), which matches closely with F-corrected estimates observed in a previous study (Crnokrak and Roff 1999). Using the formula for inbreeding depression $\delta = 1 - e^{-BF}$, where $B$ is the number of gametic lethal equivalents and $F$ the inbreeding coefficient, it is straightforward to estimate the expected inbreeding depression. As we estimated inbreeding relative to the source population in Slovakia, such are also estimates of inbreeding depression. Remaining relative fitness compared to Slovak lynx was calculated as $1 - \delta$.

All translocated lynx were successfully genotyped (N=20), with the exception of two UlyCA2 females, originating from Switzerland, which both dispersed to Austria where one was illegally killed and the fate of the other is unknown.

#### 1.5 Stakeholder engagement, communication and attitude surveys

##### 1.5.1 Stakeholder engagement and communication

In anticipation of the reinforcement project, stakeholder engagement efforts were primarily directed towards increasing awareness of the precarious conservation status of the lynx population and securing public support for the translocation initiative (Marinko et al. 2020). During the translocation process, LIFE Lynx project placed significant emphasis on building continuous intensive connections with local communities and key stakeholders (i.e. hunters). This included regular communication and one-on-one meetings in the field (mostly in the frame of camera-trapping and GPS tracking of lynx), working with local school teachers, designing tourism initiatives with local providers, ongoing engagement with the media, and upheld transparent communication with the wider public through various channels, such as lectures, website, social media, publications, documentaries, and books. Special attention was given to highlight the important role played by hunters in the efforts to restore lynx population, including a documentary produced for general public about activities conducted by hunters for support of lynx recovery (https://youtu.be/fayhFYH6P68). In this way we aimed to enhance hunter stewardship for lynx conservation. Additionally, local inhabitants of the release areas were involved through “local consultative groups”, which served as platforms for constructive dialogue, addressing local concerns, and co-designing further project activities through regular meetings (Velkavrh et al. 2024).

Essential part of engaging hunters was through monitoring of the lynx population. All hunting grounds reporting lynx presence opportunistically or through their responses in questionnaires sent each year by every hunting ground in the project area were invited to participate in the survey. In Slovenia, all but one of the contacted hunting grounds responded to the invitation in the first survey year, resulting in camera trapping engaging hunters from the entire lynx distribution in the country. Moreover, the extent of the collaborating hunting grounds grew over the years, following the detected change in lynx distribution (from 36 hunting grounds in 2018-2019 to 63 in 2022-2023). Similarly, the collaboration with hunters and the extent of the survey improved over the years in Croatia. In each collaborating hunting ground, at least one hunter expressed his/her willingness to operate the camera traps during the entire survey period (September to April) on a voluntary basis. In total, more than 100 hunters participated in camera trapping during the 5-year monitoring program in Slovenia, with almost a double number of collaborators within the entire surveyed area including Croatia and Italy. The LIFE Lynx project personnel trained the hunters to operate the cameras and joined them for camera set up at the start of each survey year. The hunters suggested the optimal sites, checked the cameras on a monthly basis and retrieved the SD cards, and the LIFE Lynx personnel processed the recordings, extracted and identified the lynx and assessed the status of the lynx population on an international level. Besides meeting with the hunters to set up the camera traps and exchange the SD cards, regular feedback from the LIFE Lynx personnel was ensured via publications for hunting magazines, social media, special events and frequent personal communication with every collaborating hunter. Over the years of collaboration, we have established trustful relationships with hunters over most of the lynx distribution in the Dinaric Mountains and SE Alps. With their involvement, hunters have maintained their proactive role in lynx research and conservation, an essential aspect for safeguarding the persistence of the Dinaric SE-Alpine lynx population in the long term.

Hunters played crucial role in the translocation and releasing of lynx, as all release locations were decided in partnership with local hunters and protected area managers, who were also responsible for building the soft-release enclosures, as well as for feeding and providing security for the lynx while they were kept in the release enclosures. Furthermore, hunting organization (Hunters Association of Slovenia) lead the education of law enforcement to prevent the poaching of lynx and other wildlife. This included professional training sessions for police officers with the goal of educating them about the importance of detecting, prosecuting, and sanctioning the illegal killings. Moreover, we conducted educational seminars for field personnel (foresters, game wardens and professional hunters), who are most likely to be first to detect and report a suspected illegal killing of wild animals to the police. We also produced a handbook on the investigation of wildlife poaching (Bartol et al. 2019), which was sent to all Slovenian hunters via national hunting magazine *Lovec*.

1.5.2 Questionnaire

The main tool for data collection on public attitudes (Dataset S2) was based on a questionnaire including 48 questions (available online in Velkavrh et al. 2024). The process of developing it included identification of the relevant issues to be explored where the entire LIFE Lynx project team has participated in the subsequent design and testing of the wording of the questions. The original questionnaire was designed in English and afterwards participating national teams translated it into local languages (Croatian, Italian, and Slovenian). The questionnaire included questions covering the following topics: general sentiment towards lynx, perceptions about lynx, knowledge and beliefs about lynx, opinions about different management measures and approaches, evaluation of information sources about lynx, demographic characteristics of the respondents and LIFE Lynx project visibility.

1.5.3 Target groups and sampling

With the public attitude survey, we have targeted the main stakeholder groups which are crucial for lynx conservation – the general public and local hunters in the project area. In Slovenia, a sample of potential general public respondents was obtained from the register of inhabitants – a random stratified (Alps and Dinarics) sample of adult (18 years and older) inhabitants was obtained from the national Statistical Office. The sample included first name, last name, and address of the selected potential respondent. In Italy a commercial panel sample was used and in Croatia a CATI sample. In Slovenia questionnaires were sent to the potential respondents and an envelope with prepaid return postage was included. Seven days later a reminder/thank you card was sent to increase response rate. In Croatia, the study was conducted using a CATI telephone method. In Italy questionnaires were filled online or through telephone interviews (Mavec et al. 2024).

Sample of hunters was obtained in Slovenia by sending 3-5 questionnaires to each of the local hunting organizations in the project area and asking the leaders of the hunting organizations to distribute the questionnaires among the hunters. In Croatia a CATI telephone method was carried out and additionally an online survey was shared on various social networks and portals thematically related to the areas of Lika and Gorski Kotar. In Italy a panel survey was carried out and additionally the questionnaire was also distributed by hunters themselves to increase the sample size (Mavec et al. 2024).

1.5.4 Data management and analysis

All the data was entered into an excel form. A random sample of 3% of questionnaires entered by hand was re-checked for the typing mistakes at the end. We did not find any mistakes.

#### 2 Supplementary Results

##### 2.1 Predation and kleptoparasitism

Overall, the kill rates, prey spectrum and prey consumption (Dataset S1, Tables S1 and S2) were similar to the remnant population in the Dinaric mountains studied before (Krofel et al. 2011, 2012, 2013, 2014, 2019), which confirmed their successful integration in the local ecosystems. As the lynx population grows, it will be important to address hunter concerns regarding ungulate abundance and management. The additional predation pressure that the translocated animals, as well as the growing lynx population in general, pose on the prey species, should be appropriately addressed by the ungulate management. To account for this, ungulate management plans in Slovenia were already adjusted with participation of local hunting representatives to allow higher tolerance in reaching harvest quotas for the hunting grounds with confirmed regular lynx presence (Stergar 2024).

Beside direct impact on prey, a large number of scavenger species feeding from prey remains of the translocated lynx indicated widespread trophic interactions, similar to previous studies on the remnant lynx population (Krofel and Jerina 2016; Krofel et al. 2012, 2019). Thus, the recovery of this apex predator is increasing the amount of food available to the local scavenger communities and could benefit these species. At the same time, kleptoparasitism by scavengers could reduce the lynx food intake, especially where larger scavengers like brown bears or wild boars are abundant (Duľa and Krofel 2020; Krofel and Jerina 2016; Krofel et al. 2012).

##### 2.2 Camera trapping

Out of 12 competing models, two were ranked at the top with the difference in AIC < 2 (Table S6). Out of these, the model with highest AIC weight and cumulative weight was chosen as the model with the most support (m9; Table S6).

The baseline detection rate differed between survey years, with highest values being reached in 2021-2022 survey year (Table S7). Moreover, this parameter differed between female vs. male lynx, as well as by the characteristics of a camera trap location type (marking site, road or other location types) (Table S7). It was also evident that the distance at which the females were detected from the center of their presumed home ranges was smaller (3.94 ± 0.22 km) than males (4.98 ± 0.16 km), i.e. indicating that the females have smaller home ranges, which is also confirmed by telemetry data from the region (Dataset S1).

The state space encompassed >12,000 km$^2$ each survey year and remained relatively constant among the survey years (only 6.5% larger state space in 2022-2023 survey year compared to 2019-2020). The standard error of the density and abundance estimates remained almost unchanged over the years, and the bias of the density estimates remained negligible (Efford and Mowat 2014) throughout the survey years. The transience rate ranged between 0.34 (between survey years 2019-2020, 2020-2021 and 2021-2022, 2022-2023), and 0.36 (between survey years 2020-2021, 2021-2022) (Fig. S8).

##### 2.3 Genetics

Altogether, from year 2010 on we collected and analyzed 750 genetic samples from the Dinaric Mountains and Julian Alps (Fig. S9), (hair= 301, scat=203, urine=99, saliva from prey=61, blood collected from snow=5, saliva-buccal swab=29, tissue from dead lynx=21, blood samples from captured lynx for telemetry=31). Additionally, we also included already published historical genetic data for the Dinaric lynx population (N=88) obtained through sampling of lynx hunting trophies in Slovenia and Croatia (Polanc et al. 2012; Sindičić et al. 2013).

As both the translocated lynx and their direct offspring are included in the estimates after 2019, we have a strong Wahlund and “isolate breaking” effects (Wahlund 1928) in the final traveling windows. Using expected heterozygosity to estimate inbreeding, we can better understand the effect of the reinforcement on the genetics of the population using empirical data.

According to the last period (40 most recently detected individuals in the dataset), in scenario without the reinforcement, inbreeding would remain high at around Fe=0.32 (Table S8), with inbreeding depression δ = 0.85 (i.e. the fitness of these animals is expected to be 15% of the fitness in the source population). Considering the effects of translocations in the Dinaric mountains with translocated animals and their offspring representing 22.5% of the population (reflecting that ratio in the sample), inbreeding would be around Fe=0.19 and inbreeding depression δ = 0.68, suggesting more than double increase in fitness compared to the pre-reinforcement. Considering the abundance estimates for the Dinaric part of the population and the numbers of detected offspring (see Results in the main text), we may already be beyond the 22.5% proportion of translocated and F1 lynx in the population. If we include samples from the Alps (scenario of fully connected Alpine stepping-stone and Dinaric Mountains subpopulations) and with the translocated animals and their offspring forming a large part of the population (40% in the sample), inbreeding would drop to 0.08, and fitness increase to four times compared to the pre-reinforcement (δ = 0.39, 61% of the source population fitness). While still high, the estimated inbreeding of Fe=0.08-0.19 is within the range of inbreeding observed in the Dinaric lynx during the 1980s (Sindičić et al. 2013) when the population was still expanding, and fits within the Bonn recommendations (Bonn Lynx Expert Group 2021).

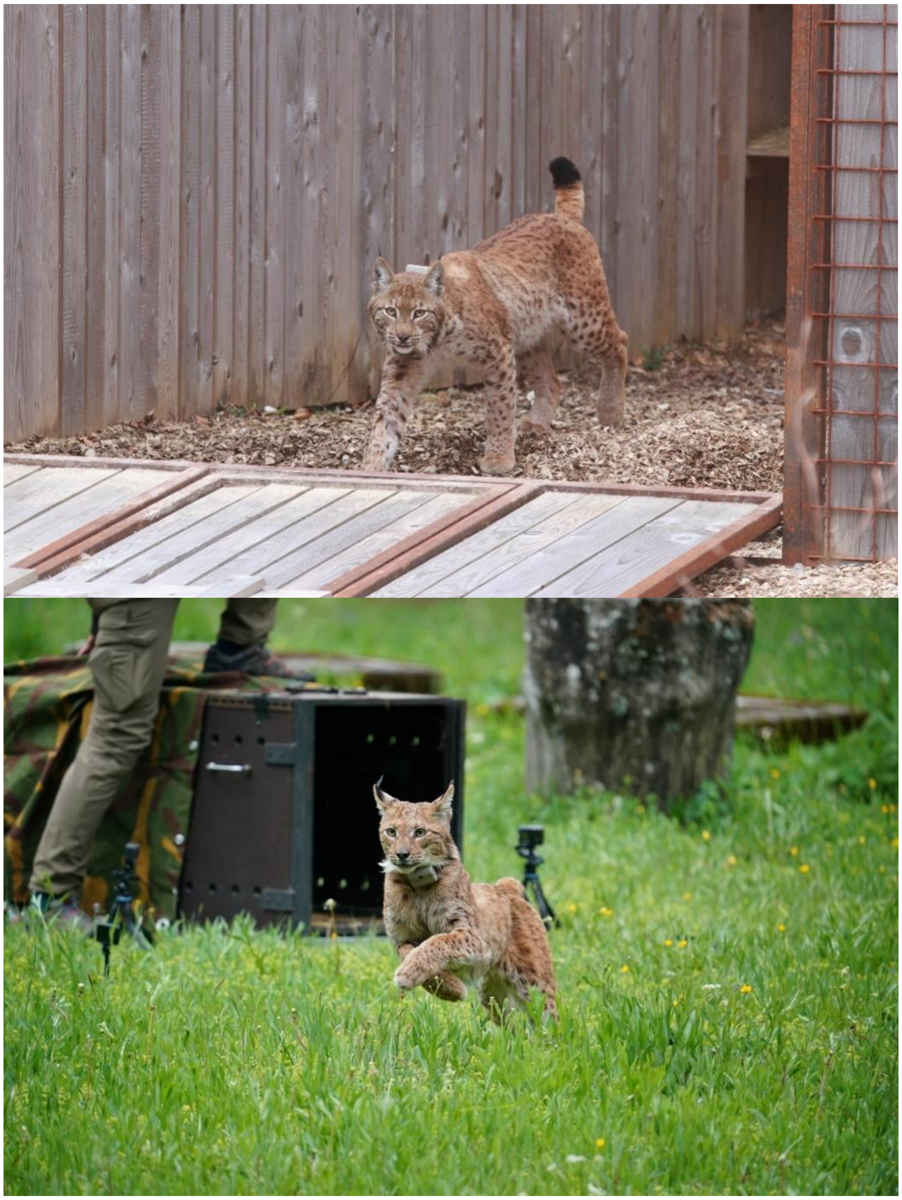

**Fig. S1.** Examples of soft-release (above, Slovenia; photo credit: M. Krofel) and hard release (below, Croatia; photo credit: M. Matešić) of lynx.

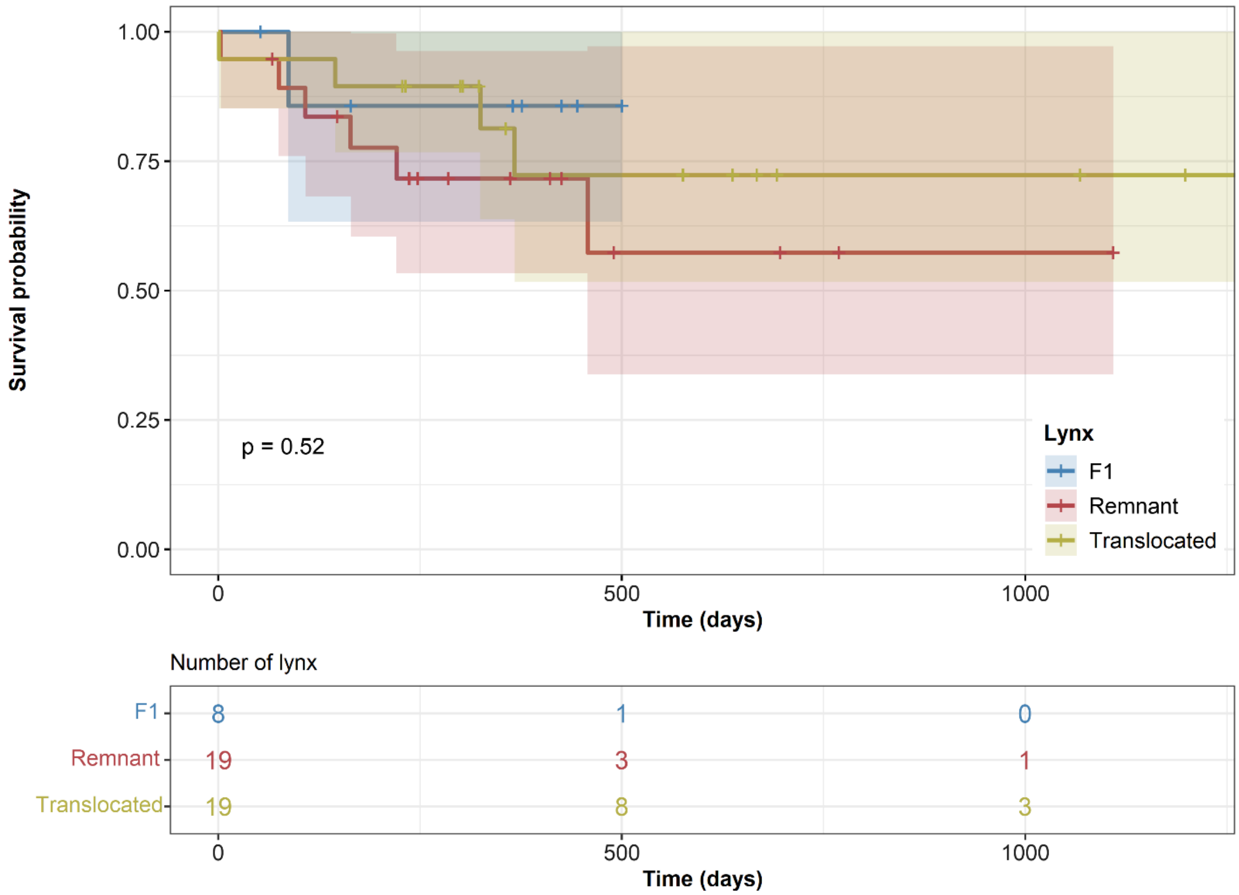


**Fig. S2.** Non-parametric survival estimates for Eurasian lynx within the tracking period. Product-limit (Kaplan-Meier) survival estimates for remnant, translocated, and F1 lynx. Lynx with unknown status at the end of the tracking period were excluded from the analysis.

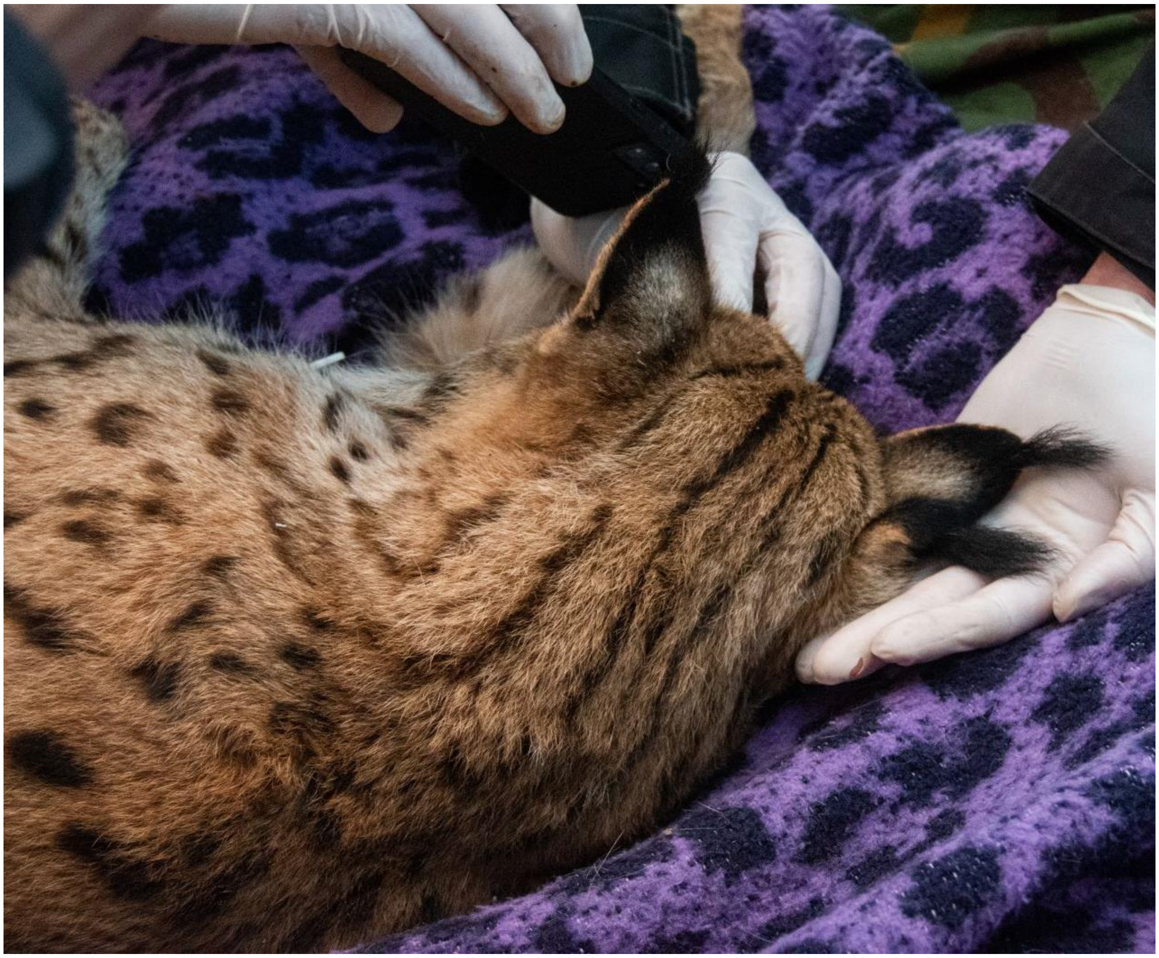

**Fig. S3.** Example of a morphological deformation in a male remnant lynx in Slovenia (double ear-tufts on the right ear), possibly associated with inbreeding (photo credit: M. Hočevar).

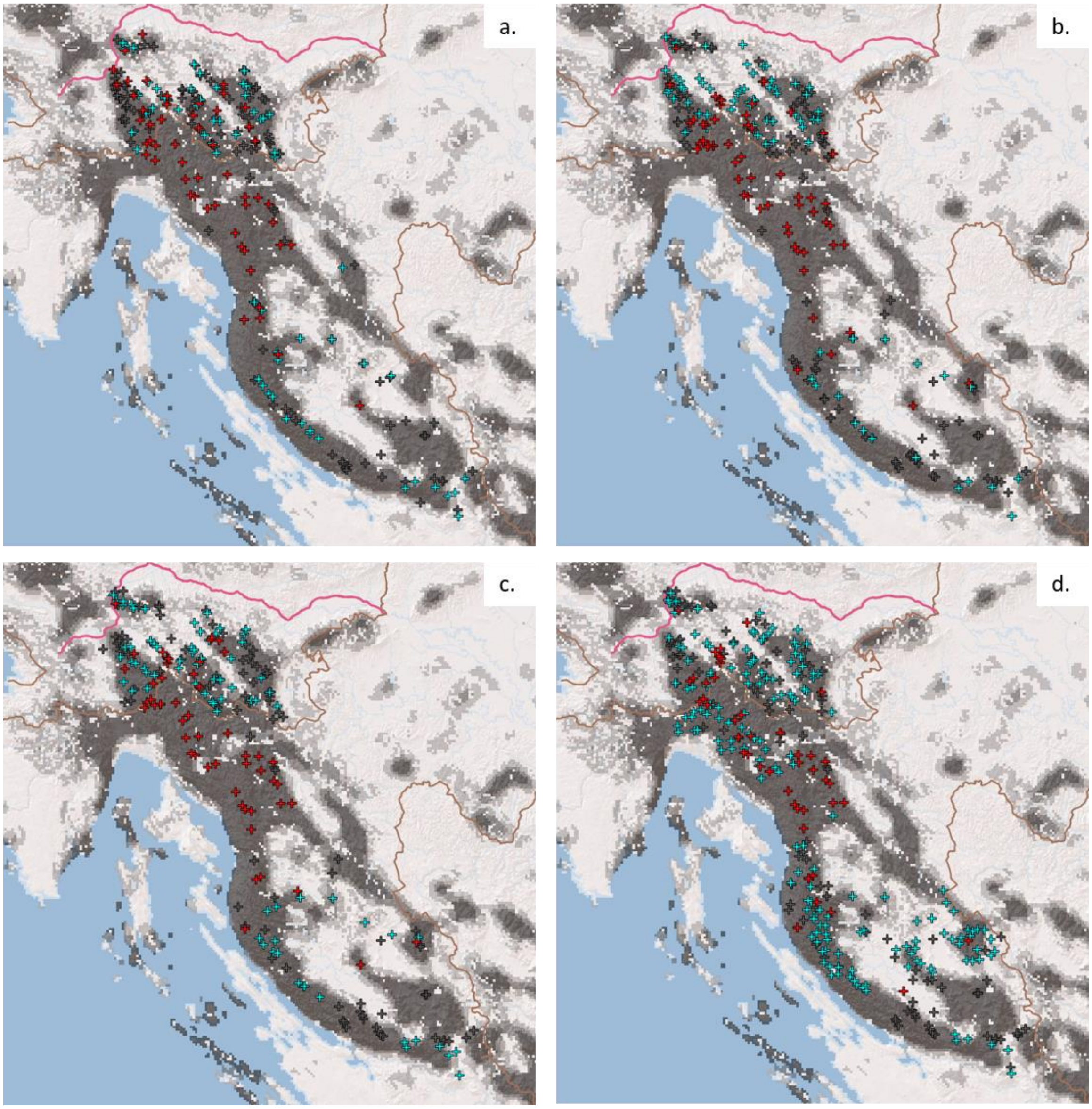


**Fig. S4.** The locations of camera traps in Northern Dinaric Mountains in each survey year (a: 2019-2020, b: 2020-2021, c: 2021-2022, d: 2022-2023). The colors of crosses show different types of location; red – marking sites, blue – roads and grey – other types of locations. Lynx habitat suitability model (Skrbinšek and Krofel 2008) is included in the maps (grey shading).

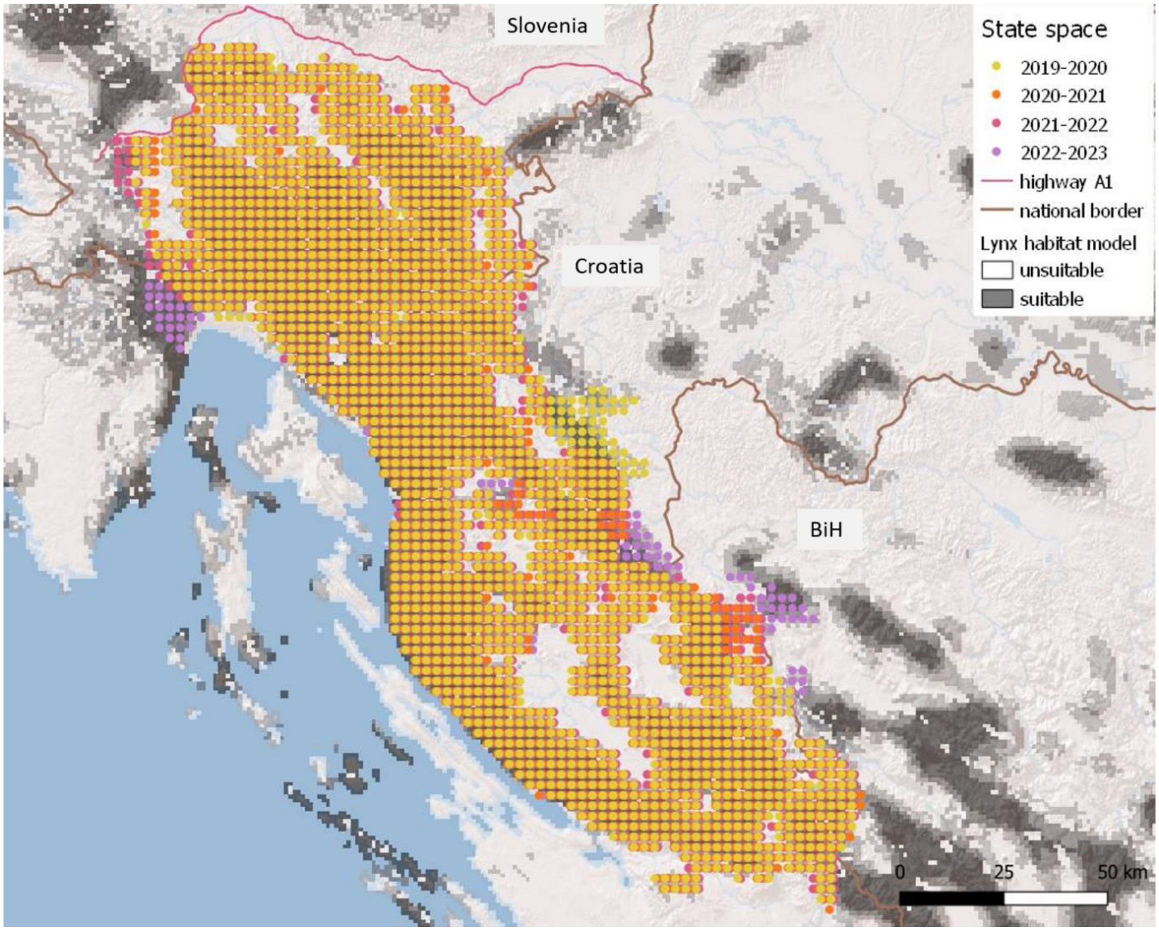


**Fig. S5.** The state space for the four survey years shown together with the habitat suitability model (Skrbinšek and Krofel 2008) and the A1 highway in Slovenia.

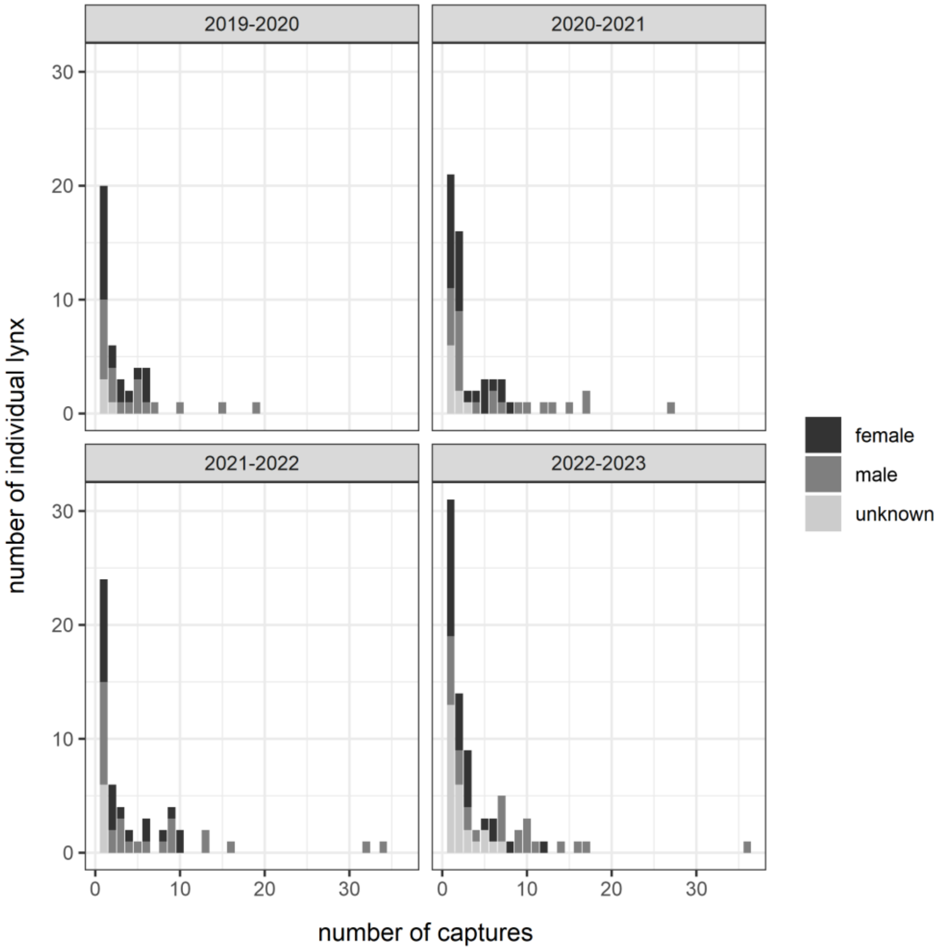


**Fig. S6.** Lynx recaptures during each survey year; the maximum number of recaptures (n=50) was recorded for a male in 2022-2023.

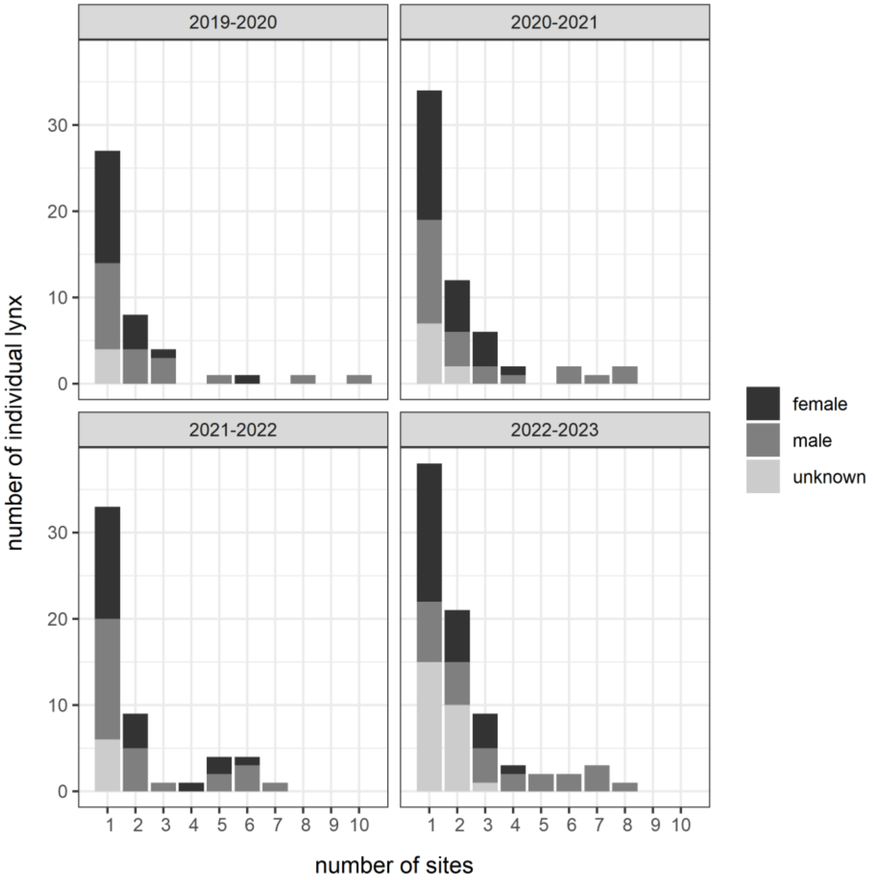


**Fig. S7.** Lynx spatial recaptures during each survey year; the maximum number of spatial recaptures (n=11) was recorded for a male in 2022-2023.

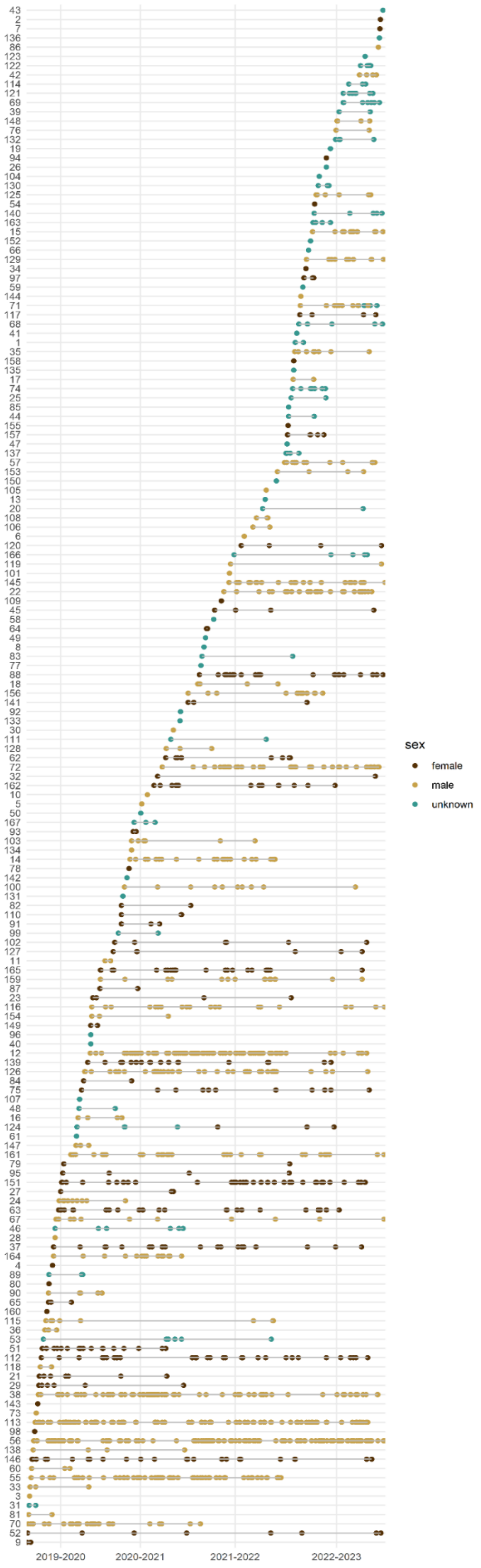


**Fig. S8.** Transience of individual lynx between the survey years. The start of each survey year is marked on the x axis while the individual lynx are shown as numbers on the y axis.

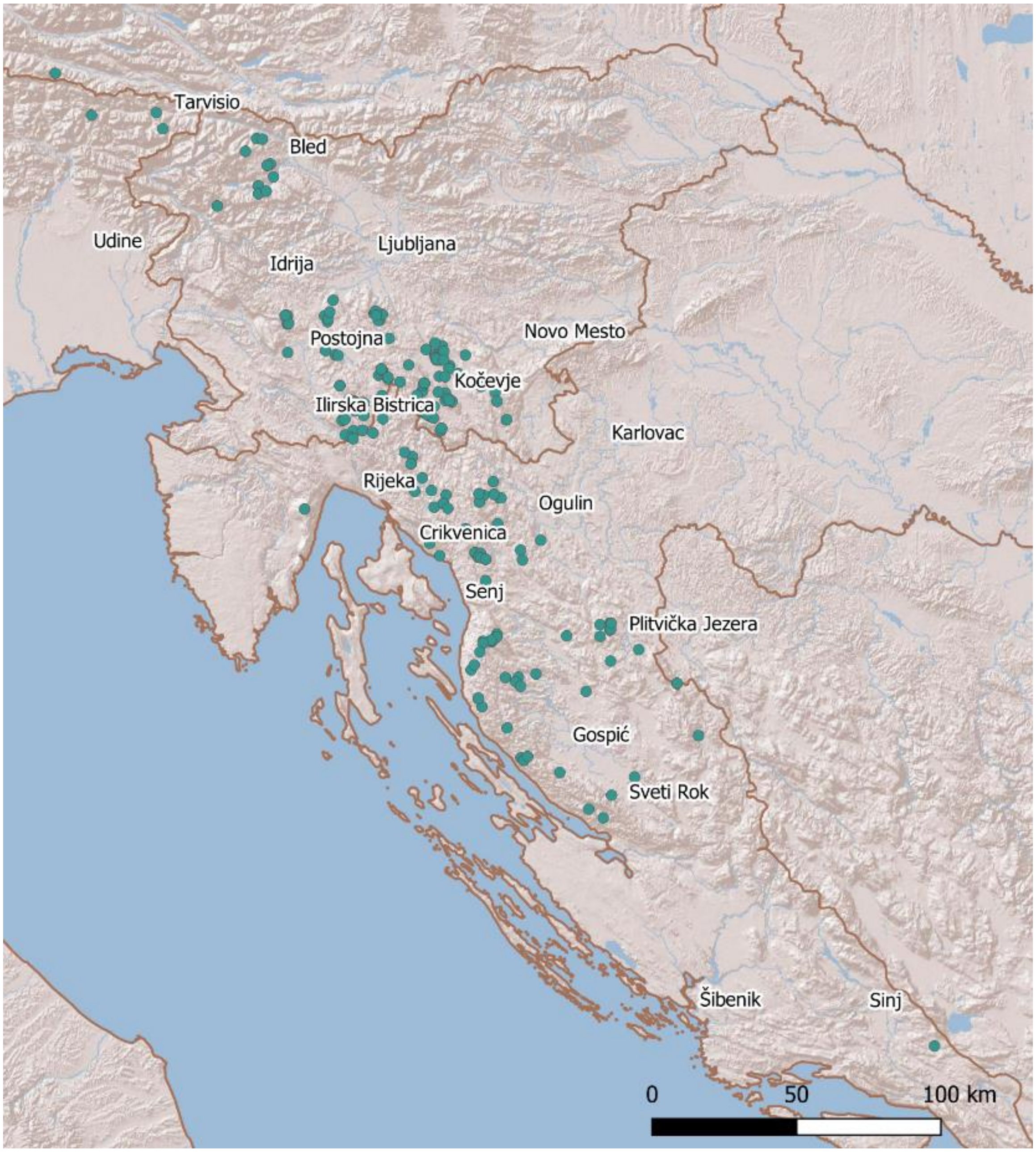


**Fig. S9.** Locations of successfully genotyped genetic samples (collected between 2010-2024), where individual lynx were identified.

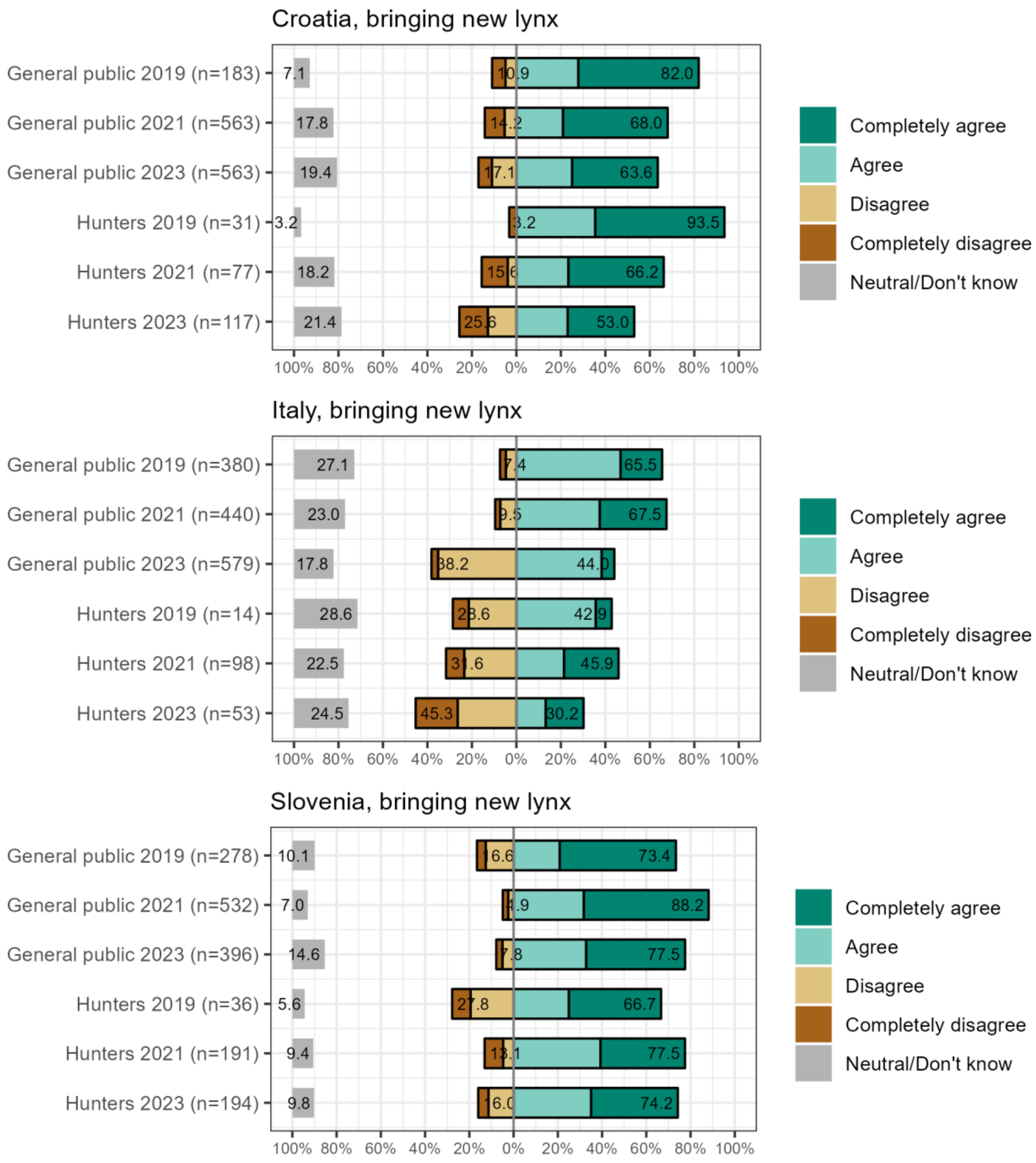


**Fig. S10.** Responses to the statement: "I support bringing new lynx to Croatia / Italy / Slovenia to save the population". Respondents answered for their respective countries.

**Table S1.** List of vertebrate scavenger species recorded feeding on kills of translocated lynx monitored with video surveillance in the Dinaric Mountains and Alps in Slovenia (n=62). Lynx were recorded as scavenging, when another lynx was recorded feeding on a kill made by translocated lynx (prey sharing).

| **Common name** | **Latin name** |
|---|---|
| Brown bear | *Ursus arctos* |
| Grey wolf | *Canis lupus* |
| Golden jackal | *Canis aureus* |
| Red fox | *Vulpes vulpes* |
| Feral dog | *Canis lupus familiaris* |
| Eurasian lynx (scavenging) | *Lynx lynx* |
| European Wildcat | *Felis silvestris* |
| Eurasian badger | *Meles meles* |
| Beech marten | *Martes foina* |
| Pine marten | *Martes martes* |
| Wildboar | *Sus scrofa* |
| Bank vole | *Myodes glareolus* |
| Golden eagle | *Aquila chrysaetos* |
| White-tailed eagle | *Haliaeetus albicilla* |
| Common buzzard | *Buteo buteo* |
| Eurasian goshawk | *Accipiter gentilis* |
| Common raven | *Corvus corax* |
| Eurasian jay | *Garrulus glandarius* |
| Coal tit | *Parus ater* |

**Table S2.** List and proportion of prey species recorded at kill sites of translocated lynx in Italy, Croatia and Slovenia (n=253).

| Common name | Latin name | Proportion among kills (%) |
|---|---|---|
| Roe deer | *Capreolus capreolus* | 83.4 |
| Red deer | *Cervus elaphus* | 8.7 |
| Chamois | *Rupicapra rupicapra* | 3.6 |
| Red fox | *Vulpes vulpes* | 1.2 |
| Mouflon | *Ovis ammon* | 0.8 |
| Brown hare | *Lepus europaeus* | 0.4 |
| European badger | *Meles meles* | 0.4 |
| European wildcat | *Felis silvestris* | 0.4 |
| Unknown | - | 1.2 |

**Table S3.** Mean female, male and total densities estimates with standard errors and 95% confidence intervals (in brackets) over four survey years for session-specific state spaces. The precision of the estimate is shown with coefficient of variation (Efford and Mowat 2014).

| | 2019-2020 | 2020-2021 | 2021-2022 | 2022-2023 |
|---|---|---|---|---|
| **population density** | 0.88 ± 0.15 (0.63-1.23) | 1.05 ± 0.15 (0.80-1.38) | 0.91 ± 0.13 (0.68-1.20) | 1.27 ± 0.15 (1.00-1.61) |
| **female density** | 0.61 ± 0.12 (0.42-0.88) | 0.72 ± 0.12 (0.52-1.00) | 0.62 ± 0.10 (0.45-0.86) | 0.87 ± 0.13 (0.65-1.16) |
| **male density** | 0.28 ± 0.05 (0.20-0.40) | 0.33 ± 0.05 (0.25-0.45) | 0.28 ± 0.05 (0.21-0.39) | 0.40 ± 0.06 (0.31-0.53) |
| **population abundance** | 110 ± 18 (79-152) | 129 ± 18 (98-169) | 112 ± 16 (84-149) | 156 ± 19 (123-198) |
| **coefficient of variation** | 0.179 | 0.156 | 0.162 | 0.141 |
| **state space ($km^2$)** | 12,350 | 12,206 | 12,350 | 12,275 |

**Table S4.** Camera trapping effort per survey year shown as i) number of camera trapping sites set on marking sites, roads and other type of locations in the Dinaric Mts., ii) cumulative number of and observed mean distance (km) between camera trapping sites (Nearest neighbor analysis tool, QGIS v. 3.6.0) per country (Slovenia, Croatia) and in total study area (Dinaric Mts) and iii) the cumulative number of camera trapping days for the Dinaric Mts.

| | | 2019-2020 | 2020-2021 | 2021-2022 | 2022-2023 |
|---|---|---|---|---|---|
| **Number of camera trapping sites** | **Marking site** | 53 | 56 | 51 | 47 |
| | **Road** | 64 | 76 | 79 | 171 |
| | **Other** | 123 | 107 | 95 | 111 |
| | **Slovenia** | 146 | 131 | 129 | 119 |
| | **Croatia** | 94 | 108 | 96 | 210 |
| | **Dinaric Mts.** | 240 | 239 | 225 | 329 |
| **Mean distance between camera trapping sites** | **Slovenia** | 1.08 | 1.29 | 1.40 | 1.47 |
| | **Croatia** | 2.34 | 2.25 | 2.52 | 1.80 |
| | **Dinaric Mts.** | 2.28 | 2.38 | 2.61 | 2.34 |
| **Number of camera trapping days** | **Dinaric Mts.** | 24,874 | 28,142 | 29,069 | 34,817 |

**Table S5.** Overview of lynx camera-trap captures for the entire study area (Dinaric Mountains) per survey year. Each survey year corresponds to a survey period which lasted from Aug 15th to Feb 15th.

| | | **2019-2020** | **2020-2021** | **2021-2022** | **2022-2023** |
|---|---|---|---|---|---|
| **Total spatial recaptures** | | 84 | 119 | 109 | 170 |
| **Total recaptures** | | 148 | 257 | 275 | 341 |
| **Mean recaptures** | | 3.44 | 4.36 | 5.19 | 4.32 |
| **Mean spatial recaptures** | | 1.95 | 2.02 | 2.06 | 2.15 |
| **MMDM** | | 10.71 | 9.14 | 10.23 | 8.63 |
| **Number of individual lynx** | *females* | 19 | 26 | 21 | 27 |
| | *males* | 20 | 24 | 26 | 26 |
| | *unknown sex* | 4 | 9 | 6 | 26 |
| | *total* | 43 | 59 | 53 | 79 |

**Table S6.** The selection of the 12 competing models. Model structure shows the effects tested per parameter, and AIC, AIC weight and cumulative weight are given for each competing model. The top-ranking models are bolded while the most supported model is additionally in italics. Session represents survey year.

| **model structure** | **model ID** | **AIC** | **AIC weight** | **Cumulative weight** |
|---|---|---|---|---|
| D~session, p0~1, sig~1 | m0 | 12978 | 4.90E-168 | 1 |
| D~session, p0~session, sig~session | m1 | 12966 | 1.80E-165 | 1 |
| D~session, p0~session+b, sig~session | m2 | 12435 | 3.00E-50 | 1 |
| D~session, p0~session+sex, sig~session | m3 | 12745 | 1.90E-117 | 1 |
| D~session, p0~session+b+sex, sig~session | m4 | 12298 | 1.50E-20 | 1 |
| D~session, p0~session+location_type, sig~session | m5 | 12831 | 2.80E-136 | 1 |
| D~session, p0~session+b+location_type, sig~session | m6 | 12360 | 8.50E-34 | 1 |
| D~session, p0~session+sex+location_type, sig~session | m7 | 12628 | 4.60E-92 | 1 |
| D~session, p0~session+b+sex+location_type, sig~session | m8 | 12226 | 7.50E-05 | 1 |
| ***D~session, p0~session+b+sex+location_type, sig~sex*** | ***m9*** | ***12210*** | ***2.80E-01*** | ***0.96*** |
| D~session, p0~session+b+sex+location_type, sig~session+sex | m10 | 12214 | 3.50E-02 | 1 |
| D~session, p0~session+b+sex+location_type, sig~1 | m11 | 12221 | 9.40E-04 | 1 |
| **D~session, p0~b+sex+location_type, sig~sex** | **m12** | **12208** | **6.80E-01** | **0.68** |

**Table S7.** Baseline detection rate across different survey years and types of camera trap location, given per lynx sex with standard error (SE) and 95% confidence interval.

| Survey year | Sex | Location type | Estimate | SE | 95% CI |
|---|---|---|---|---|---|
| 2019-2020 | f | Marking site | 0.004 | 0.001 | 0.003-0.006 |
| 2019-2020 | f | Road | 0.002 | 0.000 | 0.002-0.003 |
| 2019-2020 | f | Other | 0.002 | 0.000 | 0.001-0.002 |
| 2019-2020 | m | Marking site | 0.010 | 0.002 | 0.007-0.014 |
| 2019-2020 | m | Road | 0.006 | 0.001 | 0.004-0.008 |
| 2019-2020 | m | Other | 0.004 | 0.001 | 0.003-0.006 |
| 2020-2021 | f | Marking site | 0.005 | 0.001 | 0.003-0.006 |
| 2020-2021 | f | Road | 0.003 | 0.000 | 0.002-0.004 |
| 2020-2021 | f | Other | 0.002 | 0.000 | 0.001-0.003 |
| 2020-2021 | m | Marking site | 0.011 | 0.001 | 0.009-0.015 |
| 2020-2021 | m | Road | 0.007 | 0.001 | 0.005-0.009 |
| 2020-2021 | m | Other | 0.005 | 0.001 | 0.004-0.006 |
| 2021-2022 | f | Marking site | 0.005 | 0.001 | 0.004-0.007 |
| 2021-2022 | f | Road | 0.003 | 0.000 | 0.002-0.004 |
| 2021-2022 | f | Other | 0.002 | 0.000 | 0.002-0.003 |
| 2021-2022 | m | Marking site | 0.013 | 0.002 | 0.010-0.016 |
| 2021-2022 | m | Road | 0.008 | 0.001 | 0.006-0.010 |
| 2021-2022 | m | Other | 0.005 | 0.001 | 0.004-0.007 |
| 2022-2023 | f | Marking site | 0.004 | 0.001 | 0.003-0.006 |
| 2022-2023 | f | Road | 0.003 | 0.000 | 0.002-0.003 |
| 2022-2023 | f | Other | 0.002 | 0.000 | 0.001-0.002 |
| 2022-2023 | m | Marking site | 0.010 | 0.001 | 0.008-0.014 |
| 2022-2023 | m | Road | 0.006 | 0.001 | 0.005-0.008 |
| 2022-2023 | m | Other | 0.004 | 0.001 | 0.003-0.006 |

**Table S8.** Genetic diversity parameters estimated at the end of the translocation process (year 2024). Remnant Dinaric: calculated without the translocated lynx and their offspring, exploring situation without the effect of translocations; Dinaric w/Reinforcement: including lynx translocated to Dinaric Mountains and their offspring; Including Alp Step. St.: with all translocated lynx, including the Alpine stepping stone subpopulation. The "Reinf." Value indicates the proportion of translocated animals and their offspring in the final traveling window.

| | A | SEA | Ho | SEHo | He | SEHe | Fe | SEFe |
|---|---|---|---|---|---|---|---|---|
| Remnant Dinaric | 2.79 | 0.17 | 0.40 | 0.04 | 0.40 | 0.03 | 0.32 | 0.06 |
| Dinaric w/Reinforcement (Reinf. = 22.5%) | 3.58 | 0.30 | 0.43 | 0.04 | 0.48 | 0.03 | 0.19 | 0.05 |
| Including Alp Step. St. (Reinf. = 40%) | 3.89 | 0.35 | 0.47 | 0.03 | 0.54 | 0.03 | 0.08 | 0.04 |

**Dataset S1.** Data of GPS-collared animals (translocated lynx, F1 offspring of translocated lynx, remnant lynx) used in the study with their origin/translocation information, status and mortality causes, telemetry data, home range sizes, detected reproduction and predation in Croatia, Italy and Slovenia. We did not include lynx collared only with VHF collars. In our study we also did not consider the rehabilitated orphan lynx from the Dinaric population released in Croatia (n=2) and Italy (n=1), all of which died or disappeared before the end of the GPS-tracking period. [Dataset is currently available on request and it will be published with the peer-reviewed publication].

**Dataset S1.** Data of responses to questionnaire including 48 questions used to measure public attitudes in Croatia, Italy and Slovenia. [Dataset is currently available on request and it will be published with the peer-reviewed publication].
